# Haplotype-resolved inversion landscape reveals hotspots of mutational recurrence associated with genomic disorders

**DOI:** 10.1101/2021.12.20.472354

**Authors:** David Porubsky, Wolfram Höps, Hufsah Ashraf, PingHsun Hsieh, Bernardo Rodriguez-Martin, Feyza Yilmaz, Jana Ebler, Pille Hallast, Flavia Angela Maria Maggiolini, William T. Harvey, Barbara Henning, Peter A. Audano, David S. Gordon, Peter Ebert, Patrick Hasenfeld, Eva Benito, Qihui Zhu, Human Genome Structural Variation Consortium (HGSVC), Charles Lee, Francesca Antonacci, Matthias Steinrücken, Christine R. Beck, Ashley D. Sanders, Tobias Marschall, Evan E. Eichler, Jan O. Korbel

## Abstract

Unlike copy number variants (CNVs), inversions remain an underexplored genetic variation class. By integrating multiple genomic technologies, we discover 729 inversions in 41 human genomes. Approximately 85% of inversions <2 kbp form by twin-priming during L1-retrotransposition; 80% of the larger inversions are balanced and affect twice as many base pairs as CNVs. Balanced inversions show an excess of common variants, and 72% are flanked by segmental duplications (SDs) or mobile elements. Since this suggests recurrence due to non-allelic homologous recombination, we developed complementary approaches to identify recurrent inversion formation. We describe 40 recurrent inversions encompassing 0.6% of the genome, showing inversion rates up to 2.7×10^-4^ per locus and generation. Recurrent inversions exhibit a sex-chromosomal bias, and significantly co-localize to the critical regions of genomic disorders. We propose that inversion recurrence results in an elevated number of heterozygous carriers and structural SD diversity, which increases mutability in the population and predisposes to disease-causing CNVs.

## Introduction

Over the last two decades, numerous studies have provided whole-genome surveys into copy number variation in the human genome, with only limited discussion on balanced inversions (Sebat et al. 2004; Iafrate et al. 2004; Redon et al. 2006; Korbel et al. 2007; Kidd et al. 2008; Handsaker et al. 2015; Sudmant et al. 2015; Abel et al. 2020; Collins et al. 2020; Ebert et al. 2021). This present knowledge gap is because inversions, which are essentially copy-neutral, are one of the most difficult-to-ascertain genetic variant classes (Antonacci et al. 2009). Large inversions (>50 kbp) are often flanked by long segmental duplications (SDs) that exceed the length of sequencing reads or library inserts (Kidd et al. 2010; Sudmant et al. 2015; Vicente-Salvador et al. 2017; Chaisson et al. 2019). These properties have limited their characterization using short read whole-genome sequencing (WGS) (Sudmant et al. 2015; Abel et al. 2020; Collins et al. 2020), largely restricting prior studies to targeted hybridization and PCR assays (Puig et al. 2020; Antonacci et al. 2009; Stefansson et al. 2005). Notwithstanding these technical issues, targeted studies have demonstrated that inversions often span hundreds of kilobase pairs (kbp), play significant roles in genome biology by suppressing recombination (Sturtevant 1917), and cause disease phenotypes when inversions directly disrupt protein-coding genes or gene regulatory regions in the human genome (Spielmann et al. 2018; Puig et al. 2015; Lakich et al. 1993).

Previously, we and others reported that in the course of primate evolution regions corresponding to disease-associated microdeletions and microduplications at chromosomal regions 15q25, 15q13.3, 16p12.2, 16p11.2, 8p23.1, Xq22, 17q21.31, as well as at the hemophilia A locus (Xq28) changed their orientation multiple times between a direct and inverted state (Lozier et al. 2002; Zody et al. 2008; Antonacci et al. 2014; Catacchio et al. 2018; Maggiolini et al. 2019; Maggiolini et al. 2020; Porubsky et al. 2020; Puig et al. 2020). Based on the presence of inverted repeats with high identity at the boundaries of these regions, we hypothesized that non-allelic homologous recombination (NAHR) increases the probability of recurrent mutations during evolution, a phenomenon that we termed “inversion toggling” (Zody et al. 2008). Notably, the formation of complex SDs at the inversion flanks may make the same regions prone to recurrent deletions and duplications in humans. However, with the exception of Williams-Beuren syndrome (Osborne et al. 2001) (WBS) and Koolen de Vries (KdV) syndrome (Koolen et al. 2006), relatively little is known regarding the relationship of inversion polymorphisms and *de novo* copy number variants (CNVs) causing genomic disorders. Osborne and colleagues were among the first to report a rare 1.5 Mbp inversion polymorphism at 7q11.3 to be significantly enriched among parental transmissions of microdeletions causing WBS (Osborne et al. 2001), while with KdV syndrome, microdeletions occur almost exclusively on the inversion haplotype (Koolen et al. 2006; Sharp et al. 2006). In the latter case, it has been shown that more complex SDs have emerged in linkage disequilibrium with the inversion, and that microdeletion breakpoints in patients map precisely to directly oriented SDs that occur exclusively on the inversion haplotype (Zody et al. 2008; Itsara et al. 2012). For other loci, the relationship has been less clear owing to technical limitations in detecting inversions. Targeted assays applied to a limited number of loci have revealed shared single-nucleotide polymorphisms (SNPs) between directly oriented and inverted haplotypes, hinting at ongoing inversion toggling in humans (Zody et al. 2008; Antonacci et al. 2009; Aguado et al. 2014; Puig et al. 2020). However, the prevalence of inversion toggling throughout the human genome is unknown because of difficulties in ascertainment and genotyping.

Here, we present a sequence-level characterization of the complete spectrum of polymorphic inversions ≥50 bp in size among 41 unrelated human genomes, using three complementary approaches: (1) single-cell template strand sequencing (Strand-seq) (Falconer et al. 2012); (2) haplotype-resolved *de novo* sequence assemblies generated from Pacific Biosciences (PacBio) high-fidelity (HiFi) and continuous long reads (Ebert et al. 2021); and (3) Bionano Genomics single-molecule optical mapping (Lam et al. 2012). Integrating the data from these three genomic platforms reveals the genome-wide landscape of polymorphic inversions from 50 base pairs to several Megabase pairs (Mbp) in size and uncovers prevalent inversion toggling in human populations. We describe 40 genomic regions that are recurrently toggling in orientation in human populations, spanning 18 Mbp (∼0.6%) of the genome, with some loci estimated to have toggled up to 19 times placing the inversions on completely different haplotypes. We estimate inversion rates, ranging from 3.4 × 10^−6^ to 2.7 × 10^−4^ per locus per generation, and present mechanistic analyses on the formation of inverted sequences, implicating SDs and mobile elements. We describe a link between inversion toggling in humans and recurrent morbid CNV formation and discover novel inversions mapping to the locations of genomic disorders. Investigation of the fine structure of several genomic disorder critical regions extending from 0.42 to 1.68 Mbp demonstrates that recurrently formed inversions are frequently accompanied by locus-specific changes in the SD architecture, which includes genomic architectures likely to correspond to pre-mutational states for recurrent morbid CNV formation. Another consequence of inversion toggling appears to be an excess in the allele frequency of inverted regions. This, notably, results in an elevated number of carriers with heterozygous inversions in the human population, which we propose further promotes recurrent morbid CNV formation. Our data represent a rich population-genetic resource for studying predisposition to a key class of genetic diseases, highlighting a hitherto underexplored form of human genetic variation.

## Results

### Multi-platform inversion discovery reveals the human polymorphic inversion landscape

#### Integrated haplotype-resolved inversion discovery

We selected a diversity panel of n=44 samples from the 1000 Genomes Project (1KG) (1000 Genomes Project Consortium et al. 2015) for inversion discovery. These included individuals with ancestry from Africa (n=13), America (n=8), East Asia (n=9), Europe (n=8), and South Asia (n=6), respectively. We performed Strand-seq in nine samples and combined these data with previously generated data from three orthogonal platforms (Strand-seq, long-read assemblies, and Bionano; **Table S1**) available for 35 samples (Chaisson et al. 2019; Ebert et al. 2021). Excluding three related family members (children in family trios), inversions were discovered in 41 unrelated individuals (82 unique haplotypes). We generated an integrated inversion callset, which, after filtering, comprises 729 inversion sites (**Table S2**). These contain different classes of sequence inversion: (*i*) 330 inversions internal to L1 mobile element insertion polymorphisms, discussed separately below; (*ii*) 292 balanced inversions, where at least one sample shows a confident balanced inversion of genomic sequence; (iii) 40 inverted duplications; (*iv*) 29 complex sites, where the inverted segment contains CNVs or where estimated inversion genotype likelihoods (**Table S3, Fig. S1**) were of low confidence (threshold for confident genotypes: likelihood ratio over reference state > 10^3^); and (*v*) 38 likely assembly errors (GRCh38) (or rare minor alleles), where all human haplotypes are inverted with respect to the reference (**Fig. 1A**). We devised a method for combining Strand-seq and long-reads to generate full-length chromosomal haplotypes, including inversions that previously posed difficulties to haplotype construction (**Fig. S2, Methods**).

**Figure 1.**
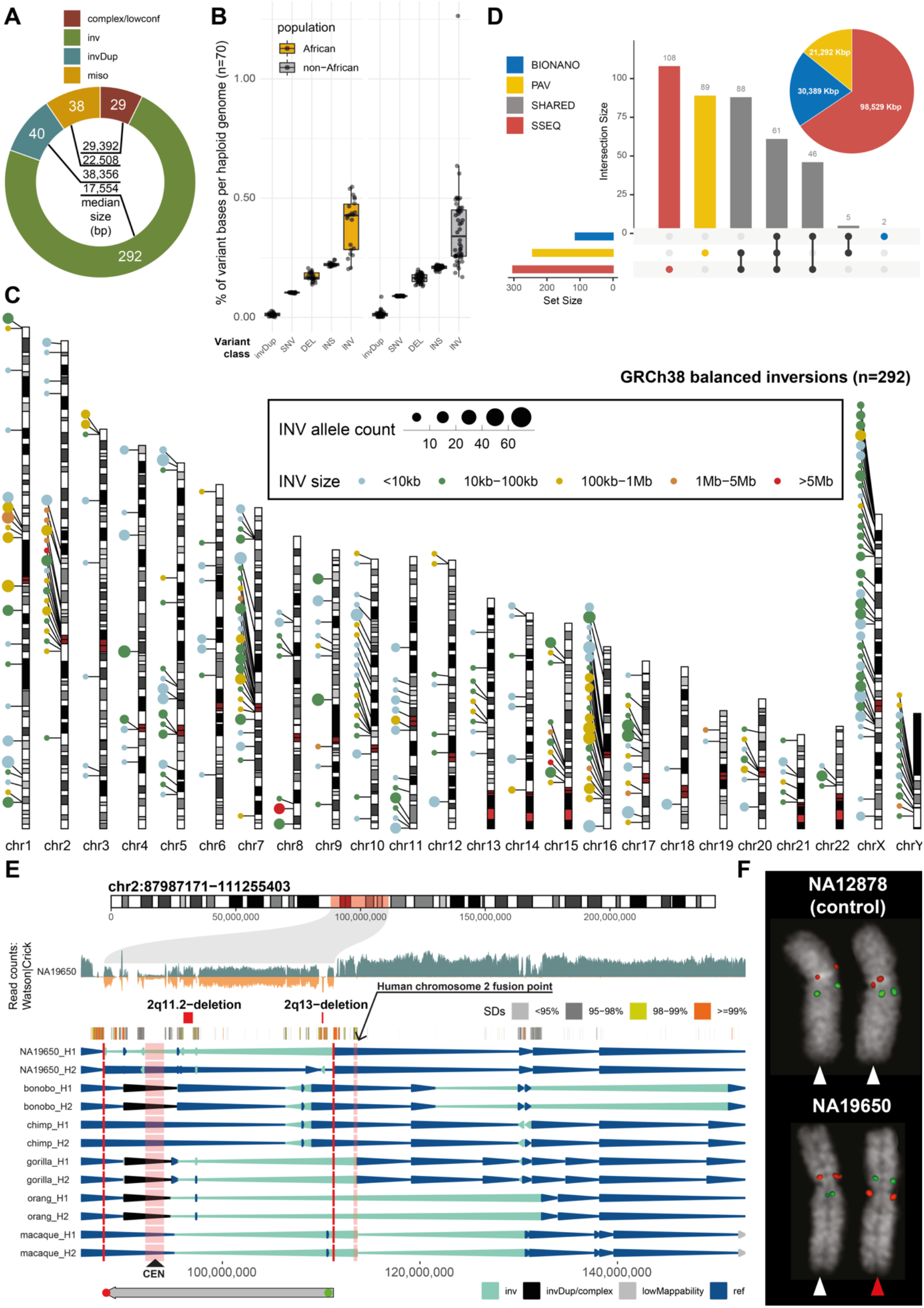
Comprehensive inversion discovery in a diversity panel. **A)** Distribution of inversion classes (L1-internal inversions presented separately in Fig. 2). **B)** Boxplot showing the percentage of affected base pairs stratified by variant class and population of origin (African and non-African). **C)** Genome-wide landscape of balanced inversions (n=292). The size of each dot represents the number of inverted alleles per inverted site across 41 unrelated samples. Color of each dot represents the size of the inversion. **D)** An upsetR plot showing the total number of inversions (n=399) independently detected by three technologies. Vertical bars report counts for technology-specific inversions (‘PAV’ - phased assembly variant caller, ‘BIONANO’ - Bionano optical maps, and ‘SSEQ’ - Strand-seq) and inversions called by at least two independent technologies (‘SHARED’). Inset: A pie chart showing the number of inverted kilobase pairs contributed by each technology. **E)** A putative novel large inversion on chromosome 2. Top: Chromosome ideogram with the inverted region highlighted by a transparent red rectangle. Below: Strand-seq read distribution with binned read counts (bin size: 50 kbp, step size: 10 kbp) represented as bars above (teal; Crick reads) and below (orange; Watson reads) the midline. The inverted region displays a distinctive distribution of Watson and Crick reads. Tracks shown below highlight regions of previously defined morbid CNVs (Coe et al. 2014) and SDs colored by increasing sequence identity. An arrowhead plot below reports phased (H1 - haplotype1, H2 - haplotype2) inversions (blue - direct orientation, cyan - inverted orientation) in the index sample (NA19650) as well as in several nonhuman primates. The bottom track depicts the positions of FISH probes used for validation. **F)** FISH validation summary for chromosome 2 inversion. Sample NA12878 was used as a control with both homologs being in a direct orientation (white arrowheads). Sample NA19650 shows one chromosome in direct orientation (white arrowhead) and one in inverted orientation (red arrowhead), therefore inverted in heterozygous state (probe ABC8-2121940H19 in red maps at chr2:88223569-88269173; probe WI2-1849B17 in green maps at chr2:110712025-110745244).

Analyzing the human inversion landscape, we find an average of 11.6 Mbp, corresponding to ∼0.39% (African: 0.43%, non-African: 0.34%) of a haploid genome, are inverted (**Fig. S3**). Comparing the currently accessible portions of the human genomes, this is four times the number of base pairs affected by SNPs (1000 Genomes Project Consortium et al. 2015) and twice the number of base pairs affected by deletion and insertion structural variants (SVs) seen in phased assemblies (Ebert et al. 2021) (**Fig. 1B**). The longest (>100 kbp) balanced inversions are most abundant on chromosomes 1, 2, 7, 10, 15, 16, 17, where large interspersed SDs have emerged during primate evolution (Marques-Bonet et al. 2009; Porubsky et al. 2020; Maggiolini et al. 2020) (**Fig. 1C, Fig. S4**). We clustered the balanced inversion breakpoints by genomic location and found 30 inversion hotspots (**Table S4**), six of which map adjacently to centromeric satellite regions (<1 Mbp). Chromosomes 1, 2, 7, 10, 16, 20, X and Y appear particularly enriched, showing 10 or more inversion breakpoints in a single hotspot (**Fig. S5**). A platform comparison shows that Strand-seq yields the largest amount of inverted base pairs, in line with its ability to discover inversions in the genome regardless of the length of the flanking repeats (Sanders et al. 2016; Chaisson et al. 2019), whereas the long-read data increases the sensitivity for events smaller than 100 kbp. Bionano is the least sensitive technique, yet provides orthogonal support for inversion discovery (**Fig. 1D, Fig. S6, Supplementary Notes**).

#### Inversion validation and orthogonal platform support

We used different methods to validate the 399 inversions outside of L1 mobile element sequences. Tests for meiotic segregation in three parent–child trios show Mendelian consistency for 247/260 (95.0%) inversion sites seen in the children, which increases to 99.5% (200/201) when considering confident inversion genotypes only (genotype-likelihood ratio over reference state > 10^3^; **Table S5**). We subjected 10 randomly selected, sequence-resolved balanced inversions (0.5 kbp–366 kbp) to PCR, successfully validating both breakpoints for 9/10 and one inversion breakpoint for the tenth event (**Fig. S7**). Using Oxford Nanopore Technologies (ONT) long-read data, we attempted to validate 202 inversion events present in three samples (HG002, HG00733 and NA19240). Though there was a clear bias for orthogonal support of small inversions, the ONT data validated inversions at 107 (∼53%) sites (**Supplementary Notes**). This is consistent with the reduced accessibility of larger inversions to long-read assembly (**Fig. 1D, Fig. S6**), because standard ONT reads still struggle to traverse across large flanking SDs. Finally, we intersected these 399 events with previously reported inversions using several orthogonal methods (Sudmant et al. 2015; Sanders et al. 2016; Audano et al. 2019; Chaisson et al. 2019; Giner-Delgado et al. 2019; Puig et al. 2020). Using a 50% reciprocal overlap, 36.3% (145/399) of our inversion callset, outside of internal L1 inversions, were reported in previous studies (**Fig. S8**, **Supplementary Methods**). Overall, 258/399 (64.7%) of these inversions, including 215/292 (73.6%) of the balanced inversions, are supported by at least one orthogonal method (**Fig. 1A**, **Fig. S9**, **Table S6**).

#### Putatively novel polymorphic inversions

We next focused on likely novel sites of inversion. Our integrated callset contains 100 putatively novel balanced inversions with none or less than 10% reciprocal overlap with events from the prior reports (Sudmant et al. 2015; Sanders et al. 2016; Audano et al. 2019; Chaisson et al. 2019; Giner-Delgado et al. 2019; Puig et al. 2020). These inversions span ∼39 Mbp of the genome (**Table S7**), and five are >1 Mbp in size, including a ∼23.2 Mbp pericentromeric inversion on chromosome 2 originating from a Mexican donor (NA19650), with its distal breakpoint lying near (∼2 Mbp) the ancestral chromosome 2 fusion point specific to humans (**Fig. 1E, Fig. S10**). We used SNP genotypes generated by PanGenie (Ebler et al. 2020) (**Methods**) to infer the presence of this inversion in the full cohort (n=3,202) of 1KG samples (Byrska-Bishop et al. 2021), based on the abundance of shared rare SNPs on the inverted haplotype. This analysis identified the mother (NA19648) of the index donor as the only additional candidate carrier for this inversion in the 1KG cohort, supporting its meiotic segregation (**Fig. S11**). We performed FISH in the two suspected carriers (NA19650 and NA19648) of the inversion, validating both (**Fig. 1F**, **Fig. S11, Methods**). We similarly searched for carriers of a large (5 Mbp) 15q11-13 inversion, which revealed four additional potential carriers in the full 1KG cohort, all of which were successfully validated by FISH (see detailed discussion further below). These data show that inversions from our discovery set can be inferred in large WGS cohorts to enable the study of disease association of inversion polymorphisms that have been inaccessible to investigation previously.

### Mechanisms for inversion formation in the human genome

#### Dominant role of NAHR in balanced inversion formation

We find that the phased sequence assemblies fully traverse the majority of balanced inversions including their breakpoints (183/292, 63%), providing an opportunity to infer underlying mutational mechanisms. Most sequence-resolved balanced inversions (132/183; 72%) show flanking inverted repeats of at least 200 bp in length (**Fig. 2A, Fig. S12**), consistent with NAHR (Bailey and Eichler 2006). This fraction is in line with prior results based on fosmid sequencing (Kidd et al. 2010) (69%) but surpasses estimates for SV insertions and deletions (Ebert et al. 2021) based on phased assembly (15–25%). Out of the 132 sequence-resolved balanced inversions likely to be mediated by NAHR, 101 (77%) showed flanking inverted SDs, whereas the remainder (23%, 31) exhibited inverted mobile element sequences (L1: n=22, and Alu: n=9). Mobile elements are known to facilitate deletion and duplication rearrangements yet are not well studied for their contributions to balanced inversion formation (Song et al. 2018). In 39% of the cases (12/31), the balanced inversion breakpoints map to full-length mobile elements (6/22 (27%) for L1 pairs, and 6/9 (67%) for Alu pairs on both flanks), whereas for the remainder (19/31) at least one inversion breakpoint maps to a truncated element. Most (21/22, 95%) inversion-flanking L1 pairs display >90% pairwise sequence identity (median: 97.2%), in sharp contrast to Alu pairs, where this is the case for only 1/9 (11%) (**Fig. S13**), suggesting that Alu-flanked inversions may form through a different rearrangement process. We also find that L1-flanked inversions are on average 8.1-fold larger (median: 5.6 kbp) than Alu-flanked inversions (685 bp; *P*< 3.8e-02, two-sided t-test). In contrast, the fully assembled SD-mediated inversions (n=101) are larger than either class of mobile element mediated event (median: 9.0 kbp), and genome-wide there is significant correlation between the size of an inversion and the length of the flanking repeat (Pearson’s correlation: R=0.67, P< 3e-16, **Fig. S14**). The propensity of balanced inversions to intersect with annotated functional elements of the genome differs between these types of balanced inversions (**Fig. 2B**) with more genes inverted through SD-mediated inversions than other types, likely due to their larger size (**Fig. 2C**). The nonrandom genomic distribution of L1 elements (biased towards AT-rich and gene-poor regions) also affects their likelihood of inverting genes (Graham and Boissinot 2006).

**Figure 2.**
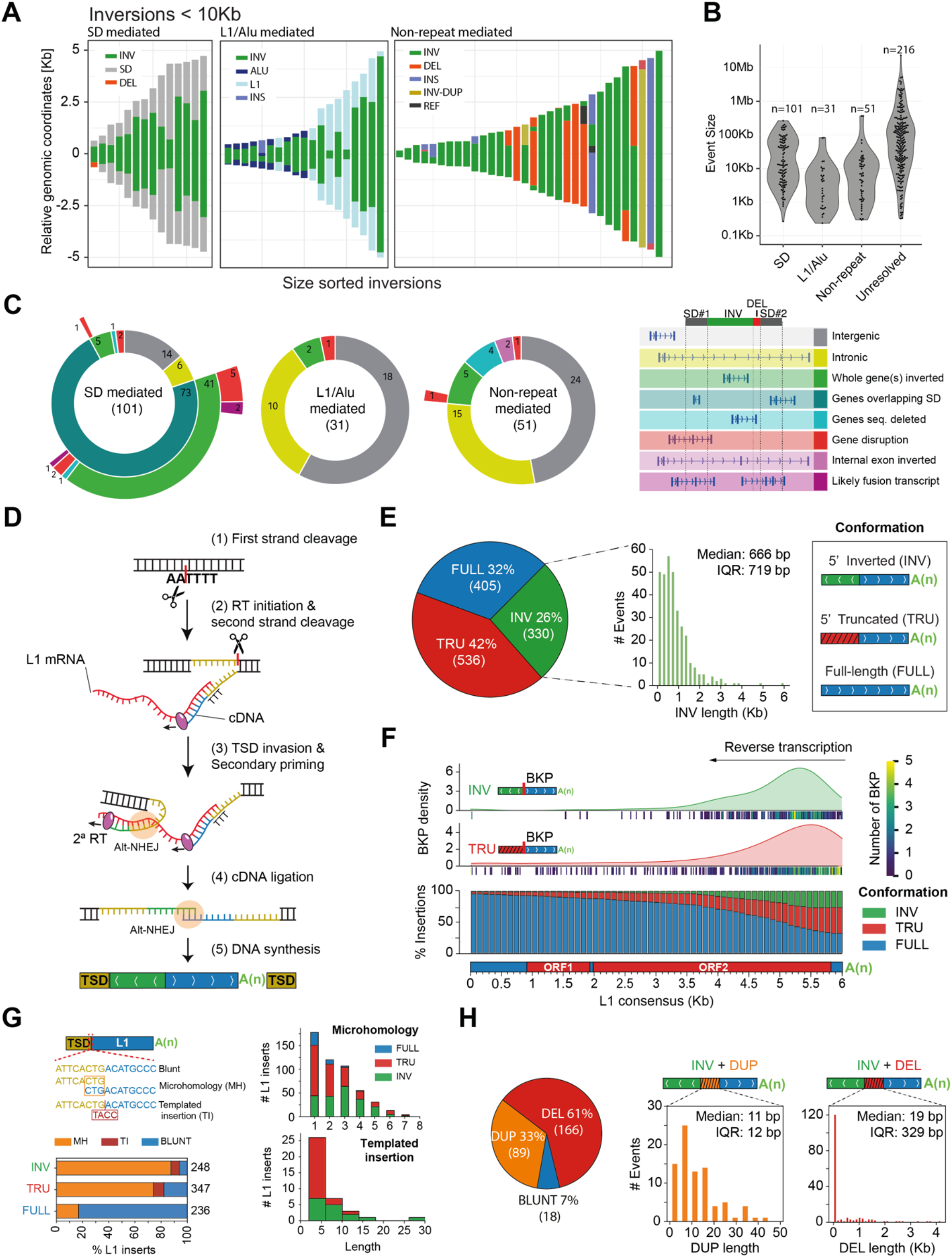
Inversion formation mechanisms. **A)** Schematic representation of inversions and their flanking sequences, displayed separately for inversions flanked by SDs (left), mobile elements (middle), and inversions not flanked by repeats (right) (here shown for events <10 kbp, and depicted for all inversions in Fig. S9). **B)** Length distribution for inversion types from panel (A). Unresolved: not fully sequence-assembled inversion. **C)** Overlap for inversion types from panel (A) with functional annotations of the genome. **D)** Mechanistic scheme depicting inversion formation at L1 insertions via twin-priming. L1 retrotransposition is initiated by the cleavage of the first DNA strand by the L1-encoded endonuclease at the target-degenerated motif of 3’- AA|TTTT-5’ (Luan et al. 1993; G. J. Cost and Boeke 1998; Gregory J. Cost et al. 2002) (1). The resulting T-rich single-stranded DNA serves for the attachment of L1 mRNA poly (A) and provides a 3’-hydroxyl that primes reverse transcription (RT) of the L1 transcript (2). After the cleavage of the second DNA strand, by a currently unknown molecular mechanism, the derived single- stranded overhang at the TSD anneals internally to the L1 transcript. We propose that this step likely occurs through Alt-NHEJ repair, serving as a primer for a secondary reverse transcription (RT) reaction (3). Then, the inverted and non-inverted cDNA products are likely ligated through Alt-NHEJ (Ostertag and Kazazian 2001) (4) and retrotransposition finalizes after the remaining DNA synthesis is completed and the second L1 DNA strand is joined to the target site by the action of a cellular ligase (5). **E)** Left, insertion conformations for polymorphic L1s. Right, inversion length distribution for twin-priming events. Interquartile range is abbreviated as IQR. **F)** Top, inversion and truncation breakpoint density, using kernel density estimation (KDE), along the L1 consensus sequence. The number of breakpoints at each genomic position is displayed as a dark blue to yellow colored gradient underneath each density distribution. The y-axis is scaled by the power of 10^-4^. Bottom, likelihood of each L1 integration outcome (INV, TRU and FULL) while L1 RT progresses towards the 5’ end of L1 mRNA sequence. **G)** Left, fraction of full-length (FULL), 5’ deleted and inverted L1 inserts exhibiting microhomologies (MH), templated nucleotide insertions (TI) and blunt joints between the 3’ end of the target site duplication (TSD) and the 5’ end of the integrated L1. Right, length distribution for microhomology and templated inversion events. **H)** Inversion junction conformations with duplicated (DUP) and deleted (DEL) pieces of L1 sequence and blunt joins. The DUP and DEL length distributions are displayed as histograms.

Of the sequence-resolved balanced inversions, 28% (51/183) lack inverted repeats at their breakpoints, even when a more lenient 50 bp cutoff is used to define breakpoint homology. Out of these, 23 inversions are accompanied by adjacent deletions (DEL) or insertions (INS) larger than 50 bp in size (**Fig. 2B, Fig. S12**), or partake in more complex events (e.g., INS-inversion-DEL- inversion-DEL-INS; **Fig. S15**). This complexity likely arose from a mutational process—possibly involving alternative nonhomologous end-joining (Alt-NHEJ), microhomology-mediated end joining (MMEJ), or microhomology-mediated break-induced replication (MMBIR) (Sudmant et al. 2015; Carvalho and Lupski 2016; Collins et al. 2020)—rather than from accumulated SVs, as we do not detect corresponding intermediate events. Collectively, our data suggest NAHR as the predominant mechanism for balanced inversion formation (Kidd et al. 2010), with a smaller fraction (at least 12% [23/183]) of inversions likely resulting from error-prone DNA repair processes leading to more complex DNA rearrangement outcomes.

#### Analysis of inversions within L1 insertions

Since L1 elements can contain inverted segments generated during retrotransposition (Ostertag and Kazazian 2001), we specifically focused on the analysis of 1,362 polymorphic L1 elements detectable in the phased assemblies (Ebert et al. 2021), to identify and characterize compound L1 structures containing inversions at their 5’ ends. These compound integrants are likely to arise as a result of twin-priming (Ostertag and Kazazian 2001), an alternative mechanism for L1 integration (**Fig. 2D**). During twin-priming, a single-stranded DNA overhang, corresponding to the 3’ end of the target site duplication (TSD) sequence (Kazazian and Moran 1998), anneals internally with the L1 mRNA, priming a secondary reverse transcription reaction, which leads to the synthesis and ligation of two cDNA products in opposite orientations. As a consequence, twin-priming events result in the integration of L1 sequences containing an internal inversion at their 5’ ends. We analyzed 93% (1,271/1,362) L1 polymorphisms (**Methods**) for twin-priming events, and find that 26% (330/1,271) of them show characteristic 5’ inverted sequences, whereas the remaining are either full-length (405) or 5’ truncated (536) (**Fig. 2E, Table S2**). The majority (88%; 291/330) of these inverted sequences are shorter than 1.5 kbp (median: 666 bp), with the observed inversions being 5 times shorter and less variable in size with respect to a random distribution of inversion lengths (*P* = 2.3e-99; Mann-Whitney U test; **Fig. S16A**). This enrichment towards short inversion events is observed irrespectively of the position of the inverted intervals relative to the consensus L1 sequence, generating a remarkable periodic pattern (**Fig. S17**). We also find a pronounced clustering of the junction positions between the inverted and non-inverted L1 fragments towards the 3’ end of the L1 consensus sequence, with 89% (292/330) of breakpoints occurring between 4,000 and 6,000 nucleotides with respect to the 5’ end (**Fig. 2F**). L1 truncation events follow a very similar breakpoint clustering pattern, and there is no significant difference in the length distribution between 5’ truncated L1s and the 3’ sense orientation ends of twin-priming events (*P*= 7e-02, Mann-Whitney U; **Fig. S16B-D**). These patterns suggest that the first 2 kbp of L1 reverse transcription are critical for establishing processive L1 reverse transcription and its successful completion, with 73% (405/552) of L1s inserting into the genome as full-length sequences once reverse transcription reaches beyond 2 kbp.

Next, we performed a detailed analysis of the junctions between the inverted L1s and their flanking genomic sequences, corresponding to the 3’ end of the target site duplication (TSD) (Kazazian and Moran 1998). We focused on the set of 269 polymorphic L1 insertions harboring internal inversions, which have been inserted into the GRCh38 reference genome. We find microhomologies of 1-9 bp in size (n=217; median 3 bp) and non-templated insertions (n=16; median 8 bp) for 87% (233/269) of these junctions (**Fig. 2G**). These data suggest that Alt-NHEJ is involved in the annealing of the target site sequence to the L1 mRNA to initiate the second reverse transcription reaction and implicates polymerase θ in the repair and potentially the insertion of the non-templated nucleotides (Chandramouly et al. 2021). In addition, we also observe signatures of Alt-NHEJ between the TSD and the 5’ end of truncated L1 inserts, similar to a prior analysis of truncated L1 elements present in the human reference genome (Zingler et al. 2005), while microhomology is rarely seen at the integration site of full-length L1s (**Fig. 2G**). Focusing on the full set of 330 L1-internal inversions, including polymorphic events present in GRCh38, we also examined the junctions between the inverted and non-inverted fragments, which were unambiguously resolved for 83% (273/330) of the events. We find an additional level of sequence complexity, with frequent short deletions (61%; 166/273) and duplications (33%; 89/273) of L1 sequence, whereas only 7% (18/273) of events are blunt-ended and devoid of deletions and duplications (**Fig. 2H**). We detect microhomologies (median 2 bp; n=190) and non-templated insertions (median 3 bp; n=27) between the inverted and non-inverted L1 fragments for altogether 81% (217/269) of the non-reference L1s (**Fig. S18**). This is consistent with previous data from a more limited number of L1 insertions (Ostertag and Kazazian 2001) and indicates that the internal ligation of inverted and non-inverted reverse transcription products likely occurs via Alt-NHEJ as well. Collectively, these sequence features suggest a major role for Alt-NHEJ in the resolution of retrotransposition intermediates (see models depicted in **Fig. 2D** and **Fig. S19**), resulting in internal inversions or truncations of L1 sequences.

### Genome-wide analysis of inversion recurrence unveils a sex chromosomal bias in mutational toggling

#### Inversion discovery saturation and excess of common polymorphisms

Focusing on the set of balanced inversions (n=292) in our integrated callset, we estimated the rate of inversion discovery with each new haplotype-phased genome assembly added. Remarkably, the rate of new inversion discovery quickly reaches an asymptote as more genomes are assessed. Currently, we estimate an increase of only 0.2% for each new human haplotype added. This is in sharp contrast to other SV forms and represents a significant ∼2.4-fold reduction in the rate of new inversion discovery when compared to insertions and deletions (Ebert et al. 2021) (*P*=1.0e-24; two-sided t-test; see orange line in **Fig. 3A**). Concomitantly, we also observe an excess of common (minor allele frequency [MAF]>5%) inversion alleles (67%) when compared to other SV classes (48%, *P*=2.6e-11, two-tailed Fisher’s exact test; **Methods**). While genomes of African ancestry are expected to exhibit the greatest genetic diversity (1000 Genomes Project Consortium et al. 2015), we also observe a saturated inversion discovery rate with the addition of new African haplotypes (**Fig. 3A**). Hence, surprisingly, common balanced inversions in Africans are already well represented in our human diversity panel comprising just 13 African samples. These observations suggest that newly sequenced genomes are unlikely to contribute significant numbers of new balanced polymorphic inversions without further technological advances to increase detection sensitivity in the most complex areas of the genome.

**Figure 3.**
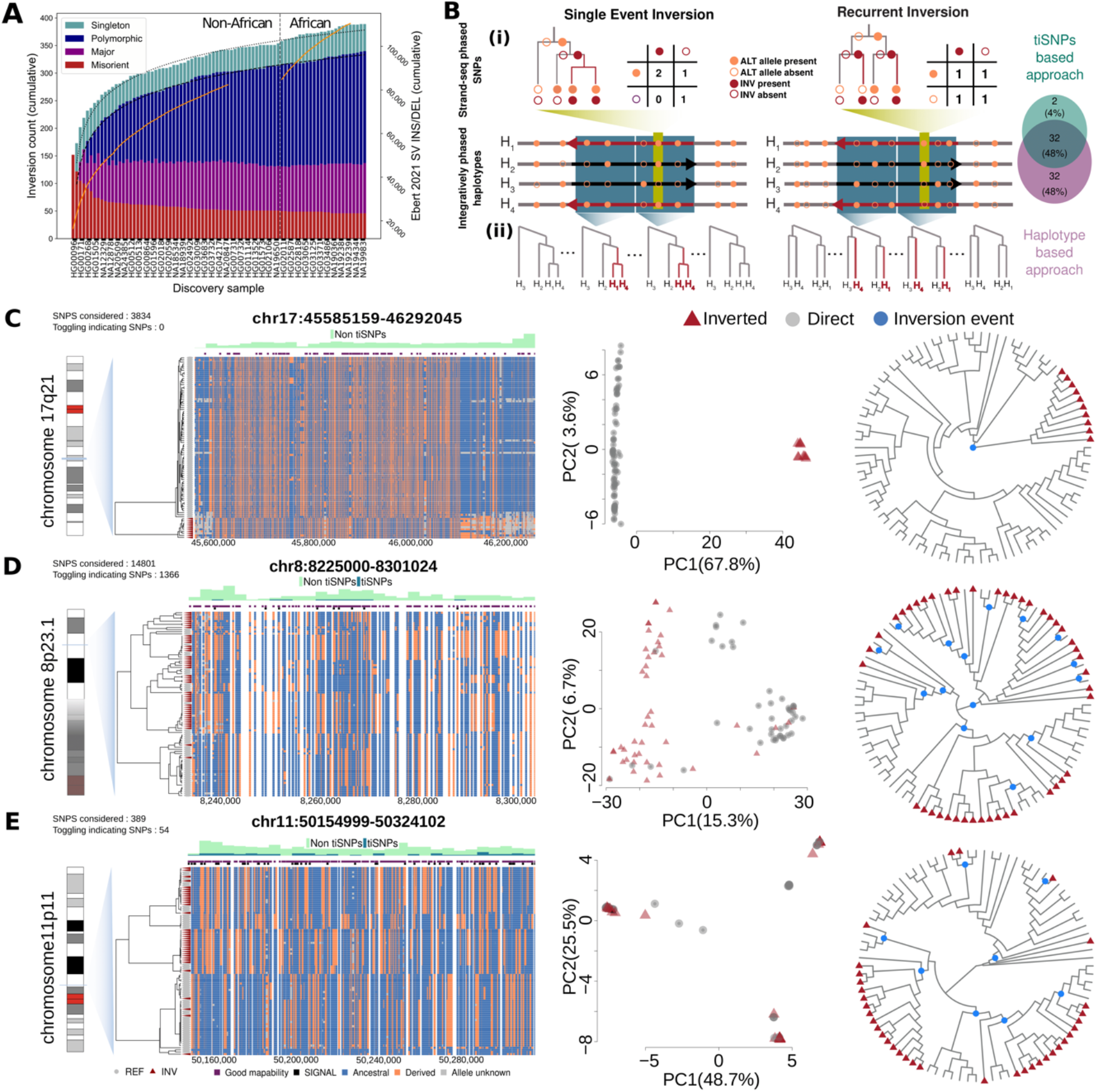
Recurrence of balanced inversions in the human genome. **A)** The rate of new balanced inversions discovered with each added genome quickly reaches an asymptote (bars and left axis). After all non-African genomes (left of the dashed line), adding new African samples increases the callset growth rate slightly (right of the dashed line). The growth rate is different from SV insertions and deletions mapped in Ebert et al. (orange line [right axis] shown for samples with assemblies in Ebert et al. 2021). Putative misorients were included to show their relative proportion. Singleton: 1 allele; Polymorphic: AF<50%; Major: AF>50% (but less than 100%), Misorient: AF=100%. **B)** Schematic representation of the two approaches devised for inversion recurrence detection. Top panel (i): tiSNPs-based approach; if an inversion occurred once in human history, there are two possible temporal orderings in which a (within inversion) SNP and the inversion could arise. In either of these cases, at most three possible SNP-inversion haplotype configurations can exist. Occurrence of each SNP allele in both inverted and non-inverted state (observing all four possible haplotype configurations) indicates that the inversion recurred. Bottom panel (ii): Haplotype-based approach that uses marginal trees inferred using Relate (v1.1.7) and IQ-Tree (v2.1.3) along with integratively phased haplotypes to evaluate individual inversions. The orientation status (red: inverted; gray: direct) of haplotypes are projected onto each marginal tree. For individual trees, monophyletic and polyphyletic groups of inverted haplotypes indicate single and recurrent inversion events, respectively. An inversion was called recurrent if 95% of the trees within the inversion locus were polyphyletic (i.e., ≥2 events; **Methods**). ALT, derived SNP alleles, determined using chimpanzee (PanTro6). Right panel: Venn diagram, showing agreement between both methods across all 127 tested inversions. **C-E)** Evidence for single (**C**, 17q21) and recurrent (**D**, 8p23.1 [distal part chr8:8225000- 8301024] and **E**, 11p11) inversion loci. The leftmost panel shows the variation of haplotypes and the relationship between inverted (red triangles) and direct (gray circles) orientations. Dendrograms of haplotypes are plotted using a centroid hierarchical clustering method. In each haplotype, ancestral and derived SNP alleles are shown in blue and orange, respectively. The histograms above the haplotypes panel show the distribution of SNPs across the locus of interest with colors distinguishing their toggling-indicating status. The positions of tiSNPs (if any) are also shown in black, in the layer immediately above the haplotype panel, and the absolute tiSNP numbers are mentioned on the top left corner. The row immediately below the histograms marks the SNPs lying in a well mappable (≥75% mappability) region. The middle panel depicts haplotype-based principal component (PC) analysis results for each locus. Inverted and direct-oriented haplotypes are shown as red triangles and gray dots, respectively. The percentage of variance explained by each PC is indicated on each axis. The right panel in each case shows the inferred cladogram of the locus of interest among sampled haplotypes using a maximum likelihood method. Red triangles are inverted haplotypes and blue dots represent the putative inversion events.

#### Methods for characterizing mutational toggling of inversions

We hypothesized that the excess of common balanced inversions may be due to recurrent mutation events flipping direct and inverted alleles back and forth due to NAHR between inverted repeats. Such “inversion toggling” in human populations would, in principle, increase allele frequency measurements for a subset of inversions. To test this hypothesis, we devised two complementary methods to infer inversion toggling (**Fig. 3B**), both of which leverage the fully haplotype-resolved nature of genetic variants in our panel. We first developed a toggling-indicating SNP (tiSNP)–based approach to identify, from the haplotype-resolved Strand-seq reads, SNPs discrepant with a single inversion origin. Discrepant biallelic SNPs were aggregated across the length of each inversion, allowing us to scan regions for signals (tiSNPs) in support for inversion toggling (**Methods**). In parallel, we developed a second haplotype-based coalescent approach to infer toggling based on the fully integrated set of phased genetic variants (generated by integrating Strand-seq and PacBio data). This approach leverages the set of internally phased SNPs along with population genetics and phylogenetic methods to find evidence in support of inversion recurrence using maximum likelihood-based phylogenetic and coalescent-based recombination graph analyses (**Methods**). The two methods are complementary as the first evaluates each SNP independently, thereby being largely unaffected by recombination, while the second leverages chromosomal linkage and variation patterns to provide confidence estimates on the number of recurrent inversion events per locus, as well as inversion rates per generation and per locus.

#### Examination of methods for detecting inversion recurrence and inversion rates

We tested and applied both methods on two previously studied inversions as controls. As a negative control, we tested the well-known 706 kbp 17q21.31 inversion (allele frequency [AF]=11%; **Fig. 3C**) that was hypothesized to have formed once in the last 2.3 million years (Koolen et al. 2006; Stefansson et al. 2005; Zody et al. 2008). As a positive control, we compared the results to the 5.3 Mbp 8p23.1 inversion (AF=50%; **Fig. 3D**) thought to be subject to a limited recurrence (Salm et al. 2012; Mohajeri et al. 2016). Using the first method we find 0% (0/3,834) and 9.2% (1,366/14,801) tiSNPs for the 17q21.31 and 8p23.1 inversion polymorphisms, respectively (**Table 1**). The tiSNPs are seen across the whole length of the 8p23.1 inversion (**Fig. 3D, Fig. S20**). In agreement with these findings, the haplotype-based coalescent approach demonstrates clear evidence for multiple recurrences of the 8p23.1 inversion at several levels, in stark contrast to the single origin of the 17q21.31 inversion. For example, a haplotype-based principal component analysis (PCA) shows that while all inverted haplotypes at 17q21.31 form a cluster distinct from the directly oriented haplotypes, the 8p23.1 locus exhibits inverted and directly oriented haplotypes appearing in similar clusters (**Fig. 3C-D**). A similar behavior is seen in hierarchical clustering-based phylogenetic trees individually constructed for the left, middle, and right one-third portions of the inversion allele (**Fig. S20**). In addition, we observe a wide distribution of identity by state among haplotypes at 8p23.1, in contrast to the distinct identity by state clusters seen for 17q21.31 haplotypes (**Fig. S21**). In total, these analyses suggest that the 8p23.1 inversion arose independently on different genetic backgrounds, in contrast to the 17q2.31 inverted haplotype which arose once and distinctly from the directly orientated haplotype.

**Table 1.**
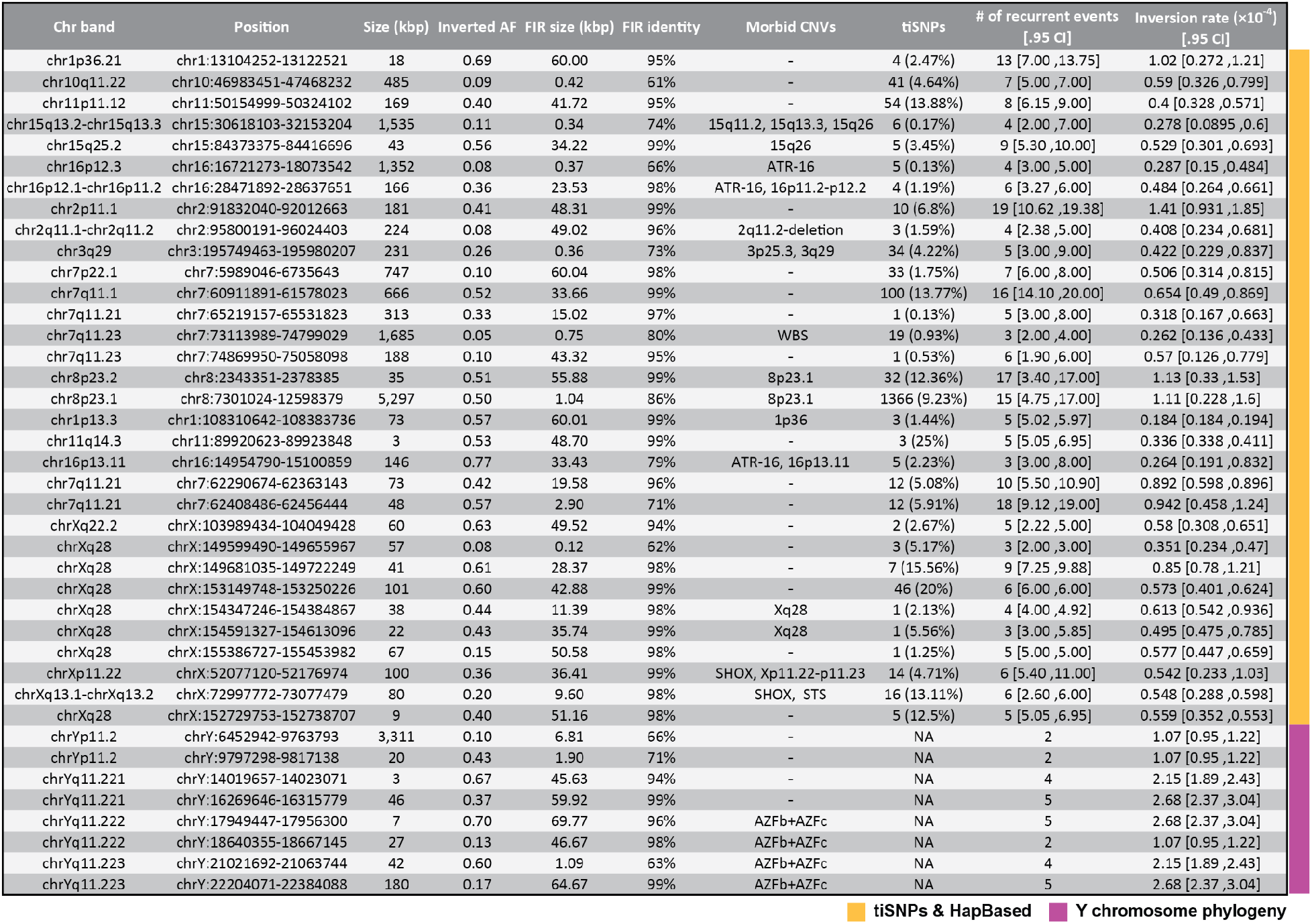
Mutational recurrence of inversions in the human genome. Abbreviations: FIR - flanking inverted repeat, AF - inversion (non-reference) allele frequency, tiSNPs - toggling- indicating SNPs, HapBased - haplotype-based coalescent approach, CI - central interval (confidence intervals are given for inversion rate estimates on the Y chromosome), WBS - Williams-Beuren syndrome. [Inversion rate estimates per generation scaled by factor 10^-4^.]

To reconstruct the evolutionary history of these balanced inversions, we inferred the underlying genealogical relationship among haplotypes (**Methods**). Because of the massive size of the 8p23.1 inversion, we focused our analysis on a 100 kbp region located at the distal portion of this locus, revealing that the inferred trees have both global bootstrap and local, marginal tree support. This suggests that the underlying genealogical relationship among haplotypes is well recapitulated (**Fig. 3D, Fig. S22**). Given the inferred tree for the 8p23.1 inversion, our method parsimoniously infers as many as 15 independent inversions occurring at this locus in humans (95% central interval: 4.75 − 17; **Methods**), with an estimated inversion rate of 1.11 × 10^−4^ per generation (95% central interval: 2.28 × 10^−5^ − 1.60 × 10^−5^). This is in contrast to the single-event 17q21.31 inversion where the same population genetics framework predicts (**Fig. 3C, Fig. S23**) an inversion rate of 3.47 × 10^−6^ (95% central interval: 2.71 × 10^−6^ − 1.03 × 10^−5^). Thus, inversion rates can vary by as much as two orders of magnitude depending on the locus.

#### Rates and genetic architecture of inversion toggling on the autosomes and the X chromosome

To understand the extent of inversion toggling in humans, we applied both the tiSNP-based approach and the haplotype-based coalescent approach to our balanced inversion callset. We focused on a subset of 127 balanced inversion sites mapping to autosomes and the X chromosome that passed a series of QC filters. Specifically, we excluded inversions in low-mappability regions, with less than 10 internal SNPs, and with ambiguous ancestral states (**Methods**). After applying these filters, we find that 52% (66/127) of inversions show evidence for inversion recurrence by at least one of the two approaches (see Venn diagram in **Fig. 3B**, **Table S8**), suggesting extensive inversion toggling in the human lineage. We constructed a “consensus“ set comprising 93 inversions based on the subset of inversions for which both approaches agree; among these, we find 32 consensus recurrent (**Table 1**) and 61 consensus single-event inversions (corresponding to an estimated fraction of 34% (32/93) toggling inversions across the autosomes and chromosome X). The remaining 34 out of 127 inversions were only called as recurrent by one of the two approaches (**Fig. 3B**), likely due to differences in sensitivity and specificity in detecting recurrence between the two approaches. Among the 93 consensus inversions, we estimate inversion rates ranging between 3.4 × 10^−6^ and 1.4 × 10^−4^ (median: 1.2 × 10^−5^). Notably, analysis of the chromosomal origin of these consensus inversions shows a significant excess of recurrent inversion loci on the X chromosome compared to the autosomes (odds ratio: 27.2, 95% C.I.: [2.55, 142.4]; *P*=1.2e-04, chi-squared test; **Fig. 4B, Fig. S24**), suggesting X-biased recurrence of inversions; however, among the consensus set of 32 recurrent inversions, we detected no significant difference in inversion rates between the X chromosome and the autosomes (*P*=0.43; Mann-Whitney U test).

**Figure 4.**
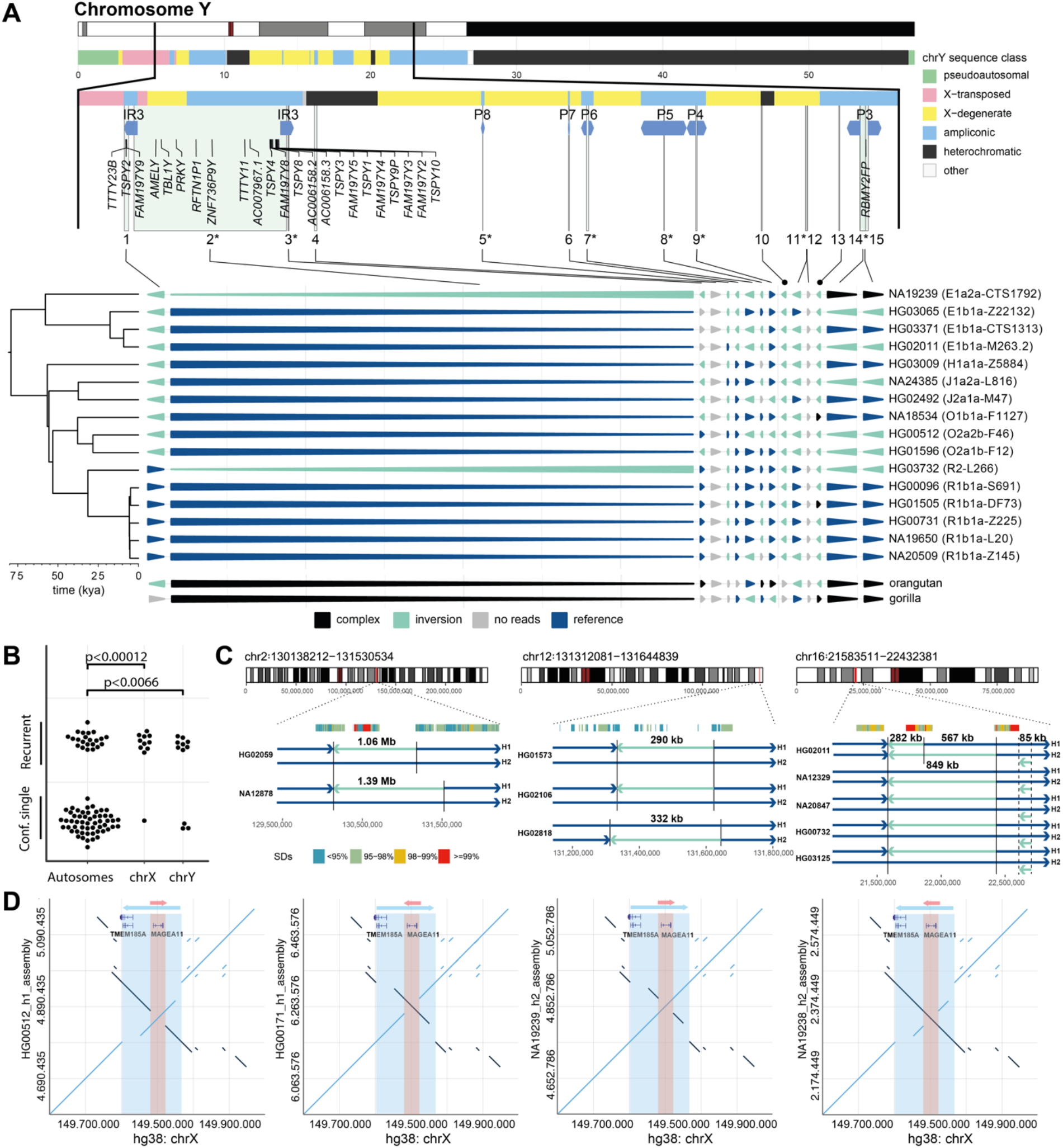
Recurrence on chromosome Y and inversion breakpoint reuse. **A)** Top: A complete chromosome Y ideogram with annotation of Y chromosomal sequence classes shown as colored rectangles. Below: A zoomed-in region (black vertical lines) of chromosome Y where all inverted sites (vertical light green rectangles) fall into. Large Y chromosomal inverted repeats flanking the called inversions (IR3 and P8-P3 as blue arrowed rectangles) and genes contained within the inverted regions are shown. Each inversion is labelled from 1-15 in the order of appearance in the arrowhead plot below, with recurrent inversions marked with an asterisk and putative misorientations/rare inversions in GRCh38 with a black dot (please note that inversions 14 and 15 are partially overlapping, the former is a ∼180 kbp recurrent balanced inversion, while the latter is a smaller ∼118 kbp inverted duplication identified only in NA19239). The arrowhead plot shows the inverted status of each region as colored arrowheads (in columns, see the legend below) per analyzed sample (in rows). The length of the arrow is proportional to the size of the inversion. Phylogenetic relationships of the analyzed Y chromosomes are shown on the left side of the arrowhead plot. **B)** Jittered dot plot showing enrichment of recurrent inversions on the sex chromosomes. **C)** Regions (n=3) with disparate inversion breakpoint reuse are highlighted on a chromosome-specific ideogram by a red rectangle. The panels below zoom into the respective regions: Top annotated SDs, colored based on sequence identity. Samples with detected inversions are shown as sample-specific haplotypes (H1 and H2) with direct (dark blue) and inverted (bright blue) regions depicted as arrows. Vertical solid lines highlight detected inversion breakpoints. **D)** Dot plots visualizing sequence alignments between GRCh38 and *de novo* phased assemblies in the region of a nested inversion on chromosome X. The outer inversion locus is highlighted in blue, the inner in red. All four possible combinations of nested inversion (inv) states (ref/ref, ref/inv, inv/ref, inv/inv; with ref. for reference orientation) were seen, suggesting inversion recurrence.

Among the consensus set of 32 recurrent inversions, we find that six overlap a set of 23 inversions previously suggested to be recurrent in the great apes (Porubsky et al. 2020). Additionally, we detect a 38 kbp recurrent inversion on the X chromosome (AF=44%; **Table 1, Fig. S25**) encompassing the genes *FLNA* and *EMD* (Small, Iber, and Warren 1997), which was previously demonstrated as recurrently inverted over the course of eutherian mammalian evolution (Cáceres et al. 2007). We predict four independent inversion events (95% central interval:4.0 − 4.92) in the past 200,000 years of human evolution, with an inversion rate of 6.13 × 10^-5^ (95% central interval: 5.42 × 10^-5^ − 9.36 × 10^-5^). Of note, this inversion is not in Hardy-Weinberg equilibrium (Fisher’s exact test *P*= 5e-03) and shows a significantly higher than expected allele frequency in African samples (AF=87.5%; global AF=42%; *P* ≤ 1e-05; 100,000 permutations). We additionally analyzed a recurrent inversion at chromosome 11p11 in more detail. We identify 54 tiSNPs (14% of 389 detected SNPs contained in the inversion), which are uniformly distributed across the inverted region (**Fig. 3E**, **Fig. S20**). Our haplotype-based approach shows that eight independent inversion events occurred at 11p11 (95% central interval: 6.15 - 9), with an estimated inversion rate of 4.0 × 10^-5^ (95% central interval: 3.28 × 10^-5^ − 5.71 × 10^-5^).

From a mechanistic perspective, we find that both the length of the flanking inverted repeat (Pearson’s correlation: 0.51; *P*= 1.7e-07) and its sequence identity (Pearson’s correlation: 0.39; *P=* 1.3e-04) positively correlate with inversion status classified as recurrent or single event (**Fig. S26**). However, flanking inverted repeat length and repeat sequence identity are themselves strongly correlated (Pearson’s correlation: 0.63, *P*= 1e-11). A multivariate logistic regression analysis shows that the major driver for the inversion status is flanking inverted repeat length (*P*= 7.2e-03), while neither repeat sequence identity nor inversion length have any significant influence (**Fig. S27**). Furthermore, analysis of flanking sequences show that the majority (72%, 23/32) of recurrent inversions on the autosomes and chromosome X exhibited ≥10 kbp flanking inverted repeat sequences with high (≥79%) sequence identity (**Table 1**). Combined, these analyses strongly implicate NAHR of flanking inverted repeats as the primary driver for inversion recurrence in the human genome helping to explain the intimate association of higher allele frequency, recurrent inversions, and flanking inverted SDs.

#### Inversion toggling affects 6% of the Y chromosome

We next turned our attention to the Y chromosome, which does not meiotically recombine with the exception of the pseudoautosomal regions. This property has the benefit of unambiguous phylogeny, greatly facilitating recurrence analyses (**Methods**). Our diversity panel, which includes 16 male samples, comprises 15 inversions on the Y chromosome (sizes: 3.4 kbp–3.3 Mbp; median: 26.7 kbp), 8 of which fell into regions that were previously reported, or suspected, to have undergone inversion (Repping et al. 2002, 2006; Lange et al. 2009; Hallast et al. 2013; Shi, Louzada, et al. 2019; Shi, Massaia, et al. 2019). The majority of inversions (13/15; 87%) are flanked by inverted SDs. A total of 10 protein-coding genes and 14 transcribed pseudogenes overlap the inverted regions, mostly confined to the large ∼3.3 Mbp Y inversion residing between the IR3 repeats (**Fig. 4A, Fig. S28**). We focused analysis of inversion recurrence on a subset of 11 balanced inversions—excluding two inversions seen on all 16 Y chromosomes, which represent minor alleles in GRCh38 or misorientations, and excluding two events with low genotype quality exhibiting too few mapped reads (**Table S9**). Out of these 11 balanced inversions, 8 (73%) were inferred as recurrent, displaying two up to five occurrences in the tree (**Table 1**). These recurrent inversions span ∼3.6 Mbp, which corresponds to ∼6% of the Y chromosomal sequence. The relative proportion of toggling inversions compared to single-event inversions on the Y chromosome is ∼7-fold higher when compared to the autosomes (*P*= 6.6e-03, chi-squared test; **Fig. 4B**), consistent with a sex chromosomal bias for inversion recurrence.

For Y chromosomal inversions identified as recurrent, we estimate inversion rates ranging from 1.07 × 10^−4^ (95% C.I.: 0.95 × 10^−4^ to 1.22 × 10^−4^) to 2.68 × 10^-4^ (95% C.I.: 2.37 × 10^-4^ to 3.04 × 10^-4^) per father-to-son Y transmission (**Table S9**). These rates correspond to one recurrent inversion per 642 (95% C.I.: 567 - 728) father-to-son Y transmissions. In further support of our measurements, we estimated a genotype concordance of 100% for four chromosome Y inversions identified and shown to be genotypable by both Strand-seq and Bionano (**Table S9**). While there have been prior reports of recurrent inversions on the Y chromosome, our analysis is the most comprehensive chromosome-wide study of inversion toggling on the Y to date. For example, the ∼3.3 Mbp IR3/IR3 inversion was previously reported to toggle at least 12 times in recent human history, with an estimated rate of ≥2.3 × 10^-4^ per father-to-son Y transmission (Repping et al. 2006). We identified this inversion in an African (NA19239) and a Southeast Asian (HG03732) individual carrying Y lineages E1a2a1a1a-CTS1792 and R2-L266, respectively, closely related to those previously reported to be inverted (**Fig. 4A**, **Fig. S28**). Our estimated inversion rate based on two events across 16 male samples is 1.07 × 10^-4^ (95% C.I.: 0.95 × 10^-4^-1.22 × 10^-4^) per generation, which is close to the published estimate. Our analyses show particularly extensive inversion toggling among large inverted SDs with >99.9% sequence identity (also referred to as Y palindromes in the literature), elements previously thought to be prone to inversion formation (Lange et al. 2009; Repping et al. 2002). The extent and rates of inversion recurrence within these structures were previously incompletely understood. We find inversions for six of the eight Y palindromes, out of which five show mutational toggling, with two (P4; ∼190 kbp long SDs) up to five (P3, P5 and P6 – 110 kbp to 495 kbp long SDs) identified inversion recurrences (**Fig. 4A**, **Fig. S29**, **Supplementary Notes**). In summary, these analyses show that inversion toggling is widespread on the Y chromosome and identify the sex chromosomes as hotspots for inversion recurrence.

#### Imprecise breakpoint reuse in further support of inversion recurrence

Because the SDs that drive NAHR occur in large blocks with multiple substrates for unequal crossover (Antonacci et al. 2014), it should be noted that recurrent inversion formation may occasionally involve disparate breakpoints, leading to a shift in the genomic coordinates of inversions on different haplotypes. Such nested inversion events would not necessarily have been classified as recurrent by our tiSNP- and haplotype-based coalescent approaches even though the unique intervening sequence has effectively toggled. We performed breakpoint mapping using the Strand-seq read data to identify partially overlapping balanced inversion polymorphisms (**Methods**). We find three regions (2q21, 12q24 and 16p12) with distinct, albeit largely overlapping, inversions (**Fig. 4C, Table S10**), suggesting that the intervening genome segment common to each inversion pair has been subject to recurrent orientation flipping in humans. All three regions reside in SD-rich regions with different SDs flanking the inversion breakpoints, suggesting that the underlying SD architecture was disparately used resulting in the formation of distinct inversions with shifted breakpoints (**Fig. 4C, Fig. S30**). We additionally identify two nearby inversions on the X chromosome with highly correlated genotypes, which exhibit one additional inversion residing in between—again within a region rich in SDs (**Fig. S31, Fig. S32**). Manual inspection revealed that these events comprise a small (41 kbp) inversion fully embedded within a larger (165 kbp) inversion, resulting in repeated toggling of the embedded segment in humans (**Fig. 4D, Supplementary Notes**). This locus, which contains the gene *MAGEA11* implicated in cancer progression (James et al. 2013; Su et al. 2019; Jia et al. 2020), was confidently identified as toggling using our tiSNP- and haplotype-based approaches (**Table 1**) and, additionally, was subject to inversion during primate genome evolution (**Fig. S32**). These data suggest that inversion recurrence is occasionally associated with disparate, partially overlapping DNA rearrangements. As sequence assembly technologies advance and precise inversion breakpoints become defined within large SDs, we anticipate that a greater fraction of inversion toggling events will be recognized as genetically distinct.

#### Inversions and gene expression

Inversions may mediate phenotypic effects by acting as expression quantitative trait loci (eQTLs) (Puig et al. 2015). Our genome-wide set of inversion genotypes provided the opportunity to identify eQTLs associated with our inversion callset. Utilizing deep transcriptomic data available for 33/44 of the samples in our diversity panel (**Data Availability**), we performed combined eQTL mapping of all inversions with a MAF of ≥1% (**Methods**), revealing 11 inversion eQTLs at a false discovery rate (FDR) of 20%. These inversion eQTLs comprise 0.55% of inversion-gene pairs (11/2007 tested sites), compared to 0.09% (56/59,464) of SV-gene pairs previously identified (Ebert et al. 2021) for deletions and 0.04% (33/85,144) for insertions, using the same FDR cutoff. Six inversion eQTLs are associated with the common 17q21.31 inversion, previously reported as an inversion eQTL using population-level PCR-based genotyping (Giner-Delgado et al. 2019), and this includes two eQTLs where the inversion is the lead variant (*MAPK8IP1P2*, *AC126544.2*) surpassing other genetic variants in significance. Novel significant associations include inversions associated with the expression of *ATP13A2*, *OR4C6*, *MAGEH1*, and *RP11-460N20.4.2* (**Fig. S33**). The sample set per population used in this study is relatively small for eQTL mapping yet showcases the possibility of associating balanced inversion polymorphisms with gene expression phenotypes to study their functional consequences.

### Relationship of polymorphic inversions with morbid CNV regions

#### Recurrent inversions are hotspots for morbid CNV formation

Copy number variation analyses of children with developmental delay and adults with neuropsychiatric disorders have identified more than 30 regions of the human genome where recurrent microdeletion and microduplication occur recurrently in a single generation leading to dosage imbalance and disease. Such recurrent morbid CNVs have been described as genomic disorders (Cooper et al. 2011; Coe et al. 2014; Bragin et al. 2014; Lupski 1998) and there are well-known cases (e.g., KdV and WBS) where the presence of an inversion polymorphism in the parent has been considered a risk factor predisposing to microdeletion/microduplication (Koolen et al. 2006; Osborne et al. 2001; Antonacci et al. 2014). Given our comprehensive map of human inversion polymorphisms, we sought to test this association more formally by genome-wide permutation analysis (**Methods**). We find a significant co-localization between morbid CNVs associated with known genomic disorders (Cooper et al. 2011; Coe et al. 2014; Bragin et al. 2014) and balanced inversions detected in our callset (14%, 40/292, *P*= 2.9e-03, twofold excess; **Fig. 5A**). In addition to WBS and KdV, this includes some of the most well-known disease-causing morbid CNVs, including Prader-Willi/Angelman syndrome (PWAS), Smith-Magenis/Potocki-Lupski syndrome (SMPLS), as well as the 15q13 microdeletion and the 16p11.2 deletion associated with autism. Remarkably, most of this association was driven by inversions documented as recurrent. For such regions, the enrichment with genomic disorders is fivefold (31%, 10/32, *P*= 1e-04, **Fig. 5A**). This suggests a relationship between the mutational toggling of inversions in humans and the propensity of sites to undergo *de novo* disease-causing CNVs.

**Figure 5.**
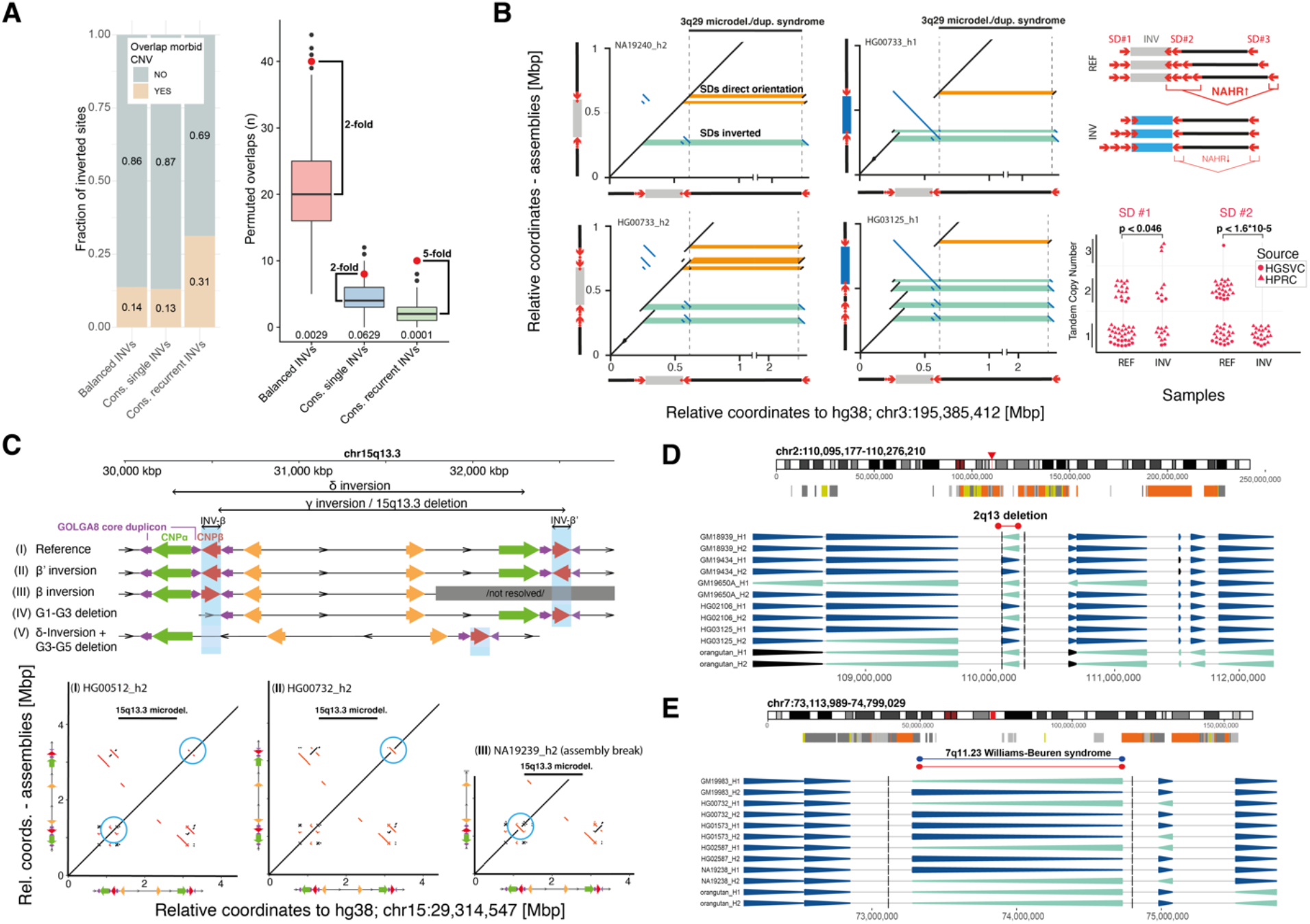
Association of toggling inversions with morbid CNV regions. **A)** Left: A barplot showing the fraction of various subsets of inversions (all balanced inversions, consensus single- event balanced inversions, and consensus recurrent balanced inversions) overlapping (allowed distance ±50 kbp) with a redundant list of morbid CNVs (n=155). Right: An enrichment analysis of overlaps between various subsets of inversions and redundant list of morbid CNVs (allowed distance ±50 kbp). The observed number of overlaps are shown by a red dot; the distributions of permuted overlap counts (n=10,000 permutations; regioneR (Gel et al. 2016) permTEST) are reported as boxplots. The bottom of each boxplot distribution provides p-values (Balanced: n=292, Consensus single: n=61, Consensus recurrent: n=32) and permuted counts. **B)** Left: Annotated dot plots of four representative assembled haplotypes at the 3q29 microdeletion/microduplication critical region. SD pairs spanning this 3q29 region and their relative orientation are highlighted in orange (direct orientation) and green (inverse orientation). One SD, contained within the inversion (top left, bottom left), exhibits an altered orientation in inversion carriers. Tandem duplications of at least one inversion mediating SD (2nd row) are observed in 43/68 (63%) of haplotypes. Right: Tandem duplications of SD#2, putatively posing a risk for morbid CNV formation through NAHR, are common in direct, but absent in inverted haplotypes (p- values based on Fisher’s exact test). HGSVC: resolved phased assemblies of the locus from our study. HPRC: open access phased assemblies obtained from the Human Pangenome Reference Consortium. **C)** Schematic view of the repeat structure found in the 15q13.3 region in different haplotypes. Three phased assembly- based dot plots illustrate structural haplotypes containing INV-β and INV-β’. Both inversions are mediated by GOLGA8 duplicons (purple arrows) and each contain a copy of the 210 kbp CNPβ repeat (red arrows), which we predict serves as a template for morbid CNV or recurrent inversion formation, depending on the combined inversion status of INV-β and INV-β’. We find additional haplotypes (IV, V) containing deletions, which are putatively protective against both inversion recurrence and morbid CNV formation (depicted in full view: Fig. S35). **D, E)** Selected regions (7q11-23, 2q13) overlapping previously defined morbid CNVs. Each plot shows a position of a given region on a chromosome-specific ideogram. Below tracks show SDs colored by increasing sequence identity and highlight the regions of previously defined morbid CNVs (Coe et al. 2014) (red - deletion, blue - duplication). The arrowhead plots below (as defined in Figure 1) report detected phased inversions in human samples and non-human primates.

#### Recurrent inversions affect the SD architecture at the 3q29 and 15q13.3 microdeletion critical regions

The propensity of inversions to intersect with morbid CNV regions led us to reassess the genomic architecture of inverted haplotypes in more detail, in order to identify haplotype-specific features that might predispose to recurrent disease-causing rearrangements. We devised a computational approach that identifies pairs of homologous SDs on the same chromosome that change their relative orientation through inversion, such that the inversion or reference (direct) orientation of a segment might represent a pre-mutational state (Zody et al. 2008) for morbid CNVs (**Methods; Fig. S34**). We find 110 balanced inversions affecting the relative orientation of altogether 1,094 SD pairs (**Table S11**), with the majority of inversions (89/110; 81%) changing the relative orientation of several (up to 112) SD pairs at once. To gain further insights we focused on those SD pairs affected by a single inversion site and considered only those sites where more than 90% of SD pairs (weighted by length) were flipped into a direct or inverse orientation, respectively—thus avoiding more complex SD regions in which inversions simultaneously rearrange a diversity of SD pairs. Based on this approach, we isolate 20 ‘potential CNV pre-mutational state inducing’ and 9 ‘potentially CNV protective’ inversions (**Table S11**).

To exemplify the utility of our approach, we characterized a recurrent inversion flanking the 3q29 microdeletion syndrome (Willatt et al. 2005; Ballif et al. 2008) critical region (AF=27%) at the sequence-level, which reorients a 21 kbp SD residing at one critical region flank (**Fig. 5B**). On the inverted haplotypes, this SD is in inverted orientation relative to the corresponding homologous SD at the distal end of the critical region, whereas non-inverted haplotypes exhibit this SD pair in a direct orientation. We further find directly oriented duplications of an SD homologous with the distal breakpoint region, which are common in directly oriented haplotypes (>50% of cases), but entirely absent in inverted haplotypes (**Fig. 5B**). These data suggest that a recurrent inversion flanking the 3q29 microdeletion critical region may act as a protective allele with respect to morbid CNV formation. Direct testing of parental carriers and transmitting haplotypes would provide proof of this hypothesis.

We further highlight the SD structure surrounding a 1.5 Mbp recurrent inversion overlapping the 15q13.3 microdeletion region, previously linked to evolutionary genome instability thought to be driven by highly identical copies of the GOLGA8 core duplicon (Antonacci et al. 2014). We find two independent inversions of approximately 210 kbp in size (denoted INV-β and β’), which encompass either copy of the CNPβ repeat previously implicated (Antonacci et al. 2014) in the formation of the 15q13.3 deletion as well as the 1.5 Mbp inversion (INV-γ) (**Fig. 5C, Fig. S35**). We hypothesize that either of the β and β’ inversions, when occurring in isolation, pose a pre-mutational state for recurrent morbid CNV formation. In contrast, when the β and β’ regions are both inverted, or when they are both in direct orientation, they may act in a protective manner towards CNV formation and may instead mediate INV-γ recurrence. We note that this model may explain why analysis of INV-β alone has not yielded a significant correlation with 15q13.3 morbid CNV formation (Antonacci et al. 2014). We also find deletions involving the CNPα and β duplicons in two haplotype structures, which potentially have protective properties both against morbid CNVs and recurrent inversions of INV-γ and δ (**Fig. S35A**, see cases IV and V), highlighting extensive structural haplotype variability at this locus.

#### Inversions at the Williams-Beuren syndrome and juvenile nephronophthisis critical regions

Encouraged by these findings, we performed more in-depth analysis of inversions intersecting sites of genomic disorders. For example, we infer that the 7q11-23 inversion (**Fig. 5D, Table 1, Fig. S36**), associated with WBS, has undergone toggling and we predict three recurrent inversion events (central interval: 2, 4) across the critical region (chr7:73,113,989-74,799,029), translating to a rate of 2.62 × 10^-5^ [central interval: 1.36 × 10^-5^, 4.33 × 10^-5^] inversion events per generation. It was previously proposed that an inversion spanning this region predisposes to morbid CNV formation (Osborne et al. 2001); however, the mutational recurrence of this inversion was previously unknown—future studies in patient cohorts may address whether a subset of these haplotypes act as a pre-mutational state for WBS. Interestingly, we additionally find a novel proximal inversion (chr7:74,984,488-75,217,887) leading to a change in orientation of a large SD, potentially resulting in an additional haplotype at risk for microdeletions at the well-known WBS critical region (**Fig. S37**).

We also observe a putatively recurrent inversion at 2q13, overlapping morbid CNVs implicated in juvenile nephronophthisis and autism (**Fig. 5E, Fig. S36**) (Parisi et al. 2004; Yasuda et al. 2014; Chen et al. 2017; Yuan et al. 2015), although recurrence of this region was inferred by the haplotype-based coalescent approach only (denoted “possibly recurrent”; **Table S8**). In this case we find two SD pairs exhibiting ≥50% reciprocal overlap with these morbid CNVs, which are predicted to change their relative orientation or architecture as a result of the inversion (**Table S11**). One of the inverted haplotypes harbors a (presumably NAHR-mediated) deletion spanning this SD pair, which may confer a protective role with respect to recurrent morbid CNV formation (**Fig. S38**). In addition, we find smaller deletions of either of the flanking SDs in several samples, which again could potentially act as protective against morbid CNV formation.

Manual investigation of HiFi-based assemblies from 28 haplotypes revealed further examples of complexity and diversity of SDs flanking common inversion polymorphisms, with the majority of such polymorphisms (11/15, 73%, Fisher’s exact test for enrichment n.s.) appearing adjacent to recurrent inversions (**Table S12**). To illustrate this breakpoint complexity, we focused on chromosome 1p36.13, which is associated with both interstitial and terminal deletions and for which we find inversion polymorphisms of the flanking SDs (**Fig. S39**) (Aagaard Nolting et al. 2020; Shapira et al. 1997). Manual analysis of optical maps of this region reveal an unprecedented level of structural diversity, with 46 distinct haplotype structures, which arose through combinations of inverted duplications, balanced inversions, and CNVs (**Fig. 6A,B**, **Fig. S40**, **Methods**). Comparison with the phased long-read assemblies reveals a high (89.6%) concordance with optical mapping data (**Methods**). The reconstructed haplotypes range between 723 kbp–1.2 Mbp in size and contain the NBPF1 core duplicon (Jiang et al. 2007). The gene *NBPF1* encodes tandemly repeated Olduvai domains (a.k.a. DUF1220), which have expanded during primate and especially human evolution potentially in association with brain size (Zimmer and Montgomery 2015; Uddin et al. 2011; Sikela and van Roy 2017; O’Bleness et al. 2012; Popesco 2006). Notably, a flanking SD that is thought to mediate morbid CNV formation (Aagaard Nolting et al. 2020) (see yellow arrow in **Fig. 6A,B**) exists in different copy number states and orientations, leaving the possibility that some of these structural haplotypes may predispose to differential susceptibility to 1p36.13 rearrangements.

**Figure 6.**
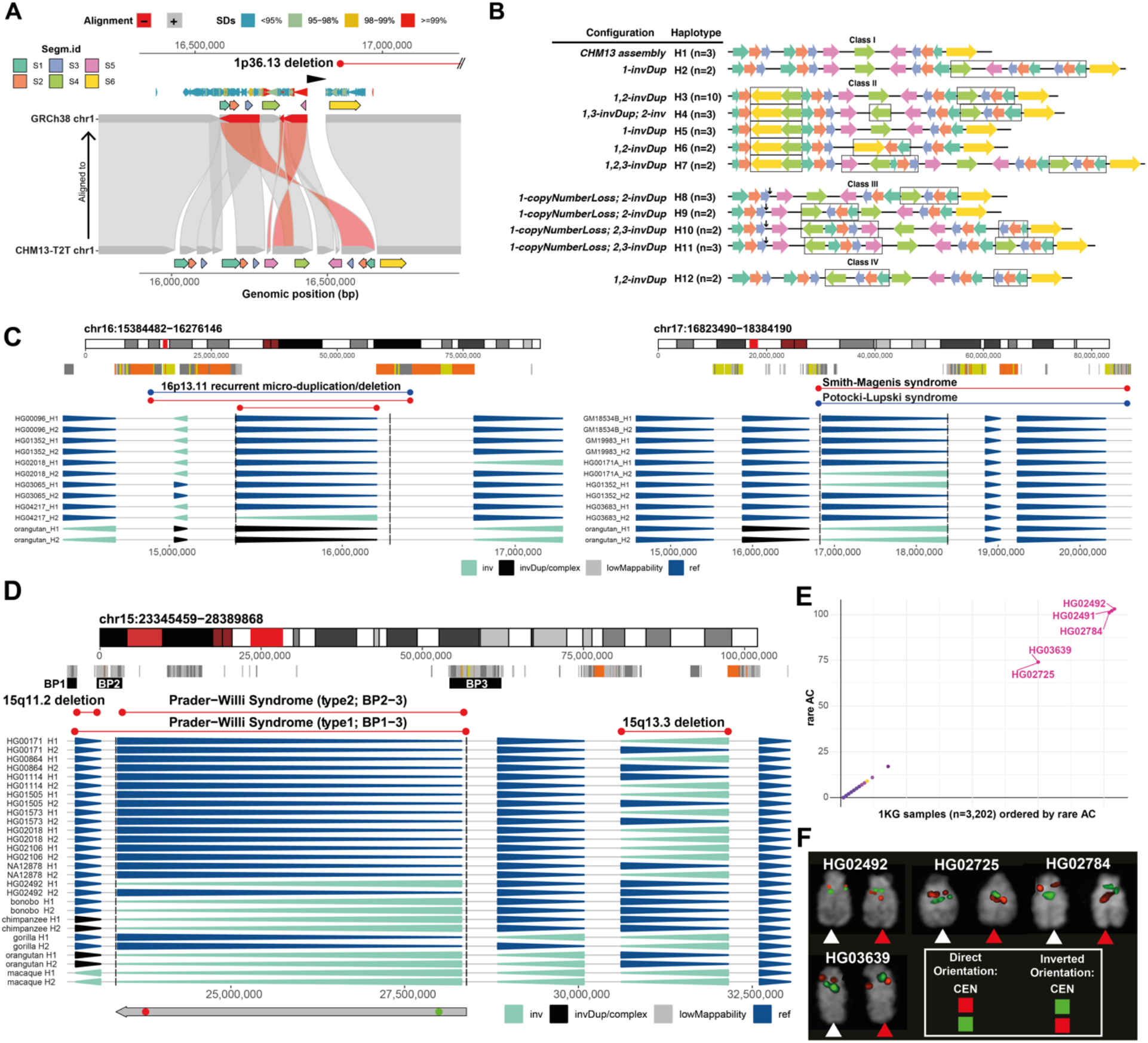
Complex inverted haplotypes and novel inversions at sites of morbid CNVs. **A)** A Miropeats-style plot highlights the structural differences between the chromosome 1p36 region of the CHM13-T2T reference genome and GRCh38 (using minimap2 alignments). Annotations provided include the alignment directionality (+ or –), SD identity, and a pathogenic CNV region at 1p36.13. **B)** Four haplotype classes consisting of 12 distinct haplotypes are seen at least twice in the 1p36.13 region, within our human diversity panel. Colored arrows represent different segments, black arrows denote deleted segments, and black rectangles represent variants observed in each haplotype compared to CHM13-T2T haplotype structure (represented as H1). The 1p36.13 haplotype structure represented in the GRCh38 reference genome was not detected in our human diversity panel. **C)** Novel inverted regions (16p13-11, 17p11-2) overlapping previously defined pathogenic CNVs. Each plot shows a position of a given region on a chromosome-specific ideogram. Below tracks show SDs colored by increasing sequence identity and highlight the regions of previously defined morbid CNVs (Coe et al. 2014) (red - deletion, blue - duplication). The arrowhead plots below (as defined in Figure 1) report phased inversions in selected human samples and non-human primates. **D)** An inversion overlapping the PWAS type II region. Arrowhead plot shown below for selected human samples and non-human primates (including the index sample HG02492). The bottom track depicts the position of FISH probes (red and green dots). **E)** A scatterplot showing the total number of shared rare inversion alleles (x- and y-axis) within the 1KG sample set (n=3,202 samples). **F)** FISH validation summary for chromosome 15 inversion in (D). Samples HG02492, HG02725, HG02784, and HG03639 show one chromosome in direct orientation (white arrowhead) and one in inverted orientation (red arrowhead) and are verified inverted in a heterozygous state.

Our analysis also discovered novel inversions that overlap well-known microdeletion syndrome critical regions not previously known to be polymorphically inverted (**Fig. 6C, Fig. S36**). This includes an inversion corresponding to the critical region of the 16p13-11 microduplication and microdeletion syndrome, which we detected in a single individual of South Asian ancestry (Indian Telugu (ITU), HG04217). SD blocks at the predicted inversion breakpoint exhibit an increased copy number in this individual in comparison to other samples in our cohort; however, this copy number increase was also seen in a sample (HG02587) lacking the inversion (**Fig. S41**). We additionally find two inversions proximal to the 16p13-11 critical region resulting in the reorientation of SD pairs, in one case (chr16:14,954,790-15,100,859, AF = 77.9%) reorienting an SD pair showing >90% reciprocal overlap with this 16p13-11 critical region, which leaves the possibility that the inversion represents a protective state for 16p13-11 morbid CNVs (**Table S11**). We additionally find a novel inversion at 17p11-2 partially overlapping the well-known SMPLS region (**Fig. 6C**), in a sample of European ancestry (Finnish, HG00171) and one of American ancestry (Colombian, HG01352). Within the complex SD architecture of the region, this inversion is predicted to lead to a reorientation of mostly directly oriented SD pairs overlapping this critical region and, thus, could potentially have protective effects with respect to microdeletion/microduplication formation at this locus (**Table S11**).

Lastly, we highlight a 5 Mbp 15q11.2-13.1 inversion, identified in our study in a single sample of South Asian ancestry (Punjabi, HG02492). The inversion overlaps the well-known PWAS type II critical region (Coe et al. 2014) and has been postulated to predispose to disease (Gimelli et al. 2003). This critical region shows a complex SD architecture at the flanks and underwent evolutionary inversion toggling in primates (Porubsky et al. 2020; Maggiolini et al. 2020) (**Fig. 6D, Fig. S42**). We set out to predict other samples carrying this inversion from our callset by subjecting the full 1KG panel to genotyping using PanGenie (Ebler et al. 2020). We detected four additional carriers (HG02725, HG02784, HG03639, and HG02491 – the mother of HG02492) in the Punjabi South Asian population, but not in other populations (**Fig. 6E, Fig. S43**). FISH validation experiments in these five predicted carriers verified all (5/5; 100%) (**Fig. 6F**, **Fig. S43, Methods**), suggesting a founder inversion event in the Punjabi population in association with this haplotype. Since the inversion is thought to be enriched in parents of Angelman syndrome patients (Gimelli et al, 2003) and potentially predisposing to disease-associated CNVs, such haplotype-based specificity in identifying carriers suggests that sequence-based phasing of inversion alleles could be used to identify families at-risk in the population.

## Discussion

### Extensive mutational recurrence of inversions in the human genome

Leveraging multiple genomic technologies, we present a comprehensive haplotype-resolved characterization of inversions, providing several new insights into a hitherto underexplored class of human genetic variation. Our analyses reveal mechanisms of formation and recurrence, as well as potential functional consequences of inversions. Amongst smaller inverted sequences (<2 kbp), we observe frequent inversion of L1 internal sequences occurring via twin-priming. Sequence analysis of the junctions between the L1 insertions and the target site duplication sequences suggests the involvement of Alt-NHEJ during both 5’ truncation and twin-priming, which typically involves terminating events or inversions occurring within the first 2 kbp of L1 reverse transcription, respectively. Full-length L1 insertions harbor different breakpoint signatures, supporting a model where the Alt-NHEJ repair machinery participates in the premature truncation and internal inversion of L1 sequences. Further research will be required to support the involvement of Alt-NHEJ, and in particular polymerase θ mediated end joining, in the DNA/RNA priming that potentially occurs at both the 5’ end of truncated sequences and internal sequences, generating inversions. In light of recent evidence of the reverse transcriptase function of this polymerase (Chandramouly et al. 2021), there may be internal priming and reverse transcription activities executed by alternative polymerases during target-primed reverse transcription.

With respect to balanced inversions, our study establishes that recurrent inversions are large, increase the frequency of heterozygous carriers, and are widespread in human populations. We report 32 balanced inversions that are subject to mutational recurrence on the autosomes and chromosome X, which corresponds to 34% of balanced inversions passing QC for recurrence assessment. On the Y chromosome, we report eight recurrent inversions. Genome-wide, all 40 recurrent inversions result in the recurrent toggling of 18 Mbp, or 0.6%, of the human genome sequence (**Table 1**). Inversion rate estimates range from 3.4 × 10^-6^ – 2.7 × 10^-4^ per site and per generation, making inversion toggling one of the most common mutational processes in the human genome. The toggled segments are gene-rich, often hundreds of kbp in length, and flanked by inverted SDs. The prevalence of ongoing inversion toggling identified in our study provides a new perspective on the evolutionary and population-based mutability of human haplotype structures. Among the 40 inversion toggling events that we characterized in the human population, we identified six (15%) that toggled between human and nonhuman ape species (Porubsky et al. 2020). This suggests that toggling has been a long-standing and persistent property for specific regions of the primate genome over the last 15 million years. Indeed, it is likely that the propensity for specific regions to toggle recurrently may in fact be even more ancient as reported for the emerin-filamin (*EMD*-*FLN*) inversion, which toggled at least 10 times across 16 distinct lineages during eutherian mammal evolution (Cáceres et al. 2007).

Because of the frequency of mutation and the fact that toggling inversions are often flanked by structurally and copy number variable regions containing genes, there are important implications for human genetics and genomics. Toggling inversions are more likely to complicate, or be missed by, eQTL mapping and genome-wide association studies because they arise independently on a diversity of haplotype structures. Unless the events are classified in association with rarer SNPs (**Fig. 6**), the inversion status will not be correctly deduced from short-read data alone. Assumptions made with respect to genomic proximity to interpret patterns of long-range gene regulation in the genome should account for genomic regions subject to recurrent toggling in orientation.

### A sex chromosome bias in mutational toggling

The analysis of the genome-wide resource of toggling inversions resulting from our study (**Table 1**) led to several important observations. First, inversion recurrence shows 27-fold and 7-fold enrichment on chromosomes X and Y compared to the autosomes, respectively, with ∼90% of the quality-filtered X chromosomal inversions and ∼70% of the Y chromosomal inversions designated as recurrent. A distinguishing characteristic of the sex chromosomes is their inability to undergo inter-chromatid homologous recombination during meiosis in a male karyotypic context, outside of their pseudoautosomal regions. In this regard, it has been hypothesized that X chromosome hemizygosity and the absence of male recombination (outside the pseudoautosomal region) may be responsible for the abundance of X chromosome inversions during eutherian mammal evolution, by promoting NAHR for unpaired X chromosomes during meiosis (Cáceres et al. 2007). DNA double-strand breaks (which could result in *de novo* DNA rearrangements) occur during meiosis on both the X and Y chromosomes, albeit at a lower density than on the autosomes (Lange et al. 2016). While most of these breaks are repaired early during meiotic prophase on the autosomes, double-strand breaks persist on the unpaired X chromosome, and the recombinases Rad51 and Dmc1 remain associated with the X chromosome in an XY karyotypic context (Moens et al. 1997; Enguita-Marruedo et al. 2019). At later stages of meiosis, factors for NHEJ co-localize at the XY bivalent (Goedecke et al. 1999). Furthermore, research in yeast has demonstrated the involvement of inverted repeat sequences in generating hairpin structures that can lead to DNA double-strand breaks or ectopic recombination between regions within the hairpin (Nag and Kurst 1997; Nasar, Jankowski, and Nag 2000; Mizuno et al. 2013); these mechanisms may also be relevant to SDs in the human genome flanking inversions in an inverted orientation. Together, these observations suggest that breaks arising in nonhomologous XY regions may be repaired by inter-sister recombination or NHEJ after crossovers have been formed, i.e., at the end of meiotic prophase when the strong inter-homolog recombination bias for repair is lifted—possibly facilitating recurrent inversion formation. In this model, the male germline may be a key driver for inversion recurrence.

### Association of recurrent inversions and hotspots of disease-causing rearrangements

Notwithstanding the sex chromosome bias, 55% of toggling inversions reside on autosomes (**Table 1**). While an intimate relationship between inversion polymorphisms and some recurrent morbid CNVs has been known for decades (Osborne et al. 2001; Koolen et al. 2006; Antonacci et al. 2009, 2014; Maggiolini et al. 2019), we demonstrate that recurrent inversions are driving this association, where the key common feature appears to be the presence of flanking large SDs. We find that toggling inversions are fivefold enriched for hotspots of morbid CNV formation. We report inversions embedded in large SDs that overlap the critical regions of a variety of microdeletion and microduplication regions, including 3q29, SMPLS, 2q13 and 15q13.3. We describe three inversions at the 16p13-11 critical region, show that inversions overlapping the WBS critical region have arisen recurrently, and identify an inversion overlapping the PWAS type II critical region as a potential founder event in the Punjabi population.

The findings from our study provide complementary mechanistic explanations for the association of inversions and hotspots of morbid CNVs: As discussed above, mutational recurrence appears to result in a remarkable abundance of high allele frequency inversions. Theoretically, the allele frequency for a recurrent inversion would converge to 0.5 if other evolutionary forces were absent, and, in this scenario, around half of all individuals in a population would be heterozygous for this inversion. In a heterozygous inverted state, homologous recombination is suppressed (Sturtevant 1917), which could facilitate further *de novo* DNA rearrangements. For example, the estimated allele frequency of the inverted state for the large 8p23.1 inversion, which we predict has arisen 15 times in humans, is 0.50 in our diversity panel (**Table 1**). Therefore, this region is predicted to be in a heterozygous inverted state in approximately 50% of all meioses, a state thought to mediate susceptibility to *de novo* copy number variation at 8p23.1 (Giglio et al. 2001; Giorda et al. 2007). A similar mechanism may be at play for other recurrent inversions identified in our study, where suppressed inter-homolog recombination in conjunction with an abundance of the heterozygous inverted state in the population may locally increase mutability. An alternative, not mutually exclusive, scenario may be that recurrent inversion formation itself generates SD complexity at the flanks irrespective of whether the locus exhibits a heterozygous inverted state. We observed a diversity of structural haplotypes in association with inversion recurrence, with large SDs being repositioned and reoriented; this in turn can promote NAHR and, depending on the relative orientation of the flanking SD pairs, trigger additional rounds of inversion or recurrent morbid CNVs.

The remarkable diversity of human haplotype structures with respect to the length and orientation of flanking SDs is exemplified by our sequence-level analysis of inversions affecting chromosomes 7q11.23 (WBS), 15q13, and 2q13 as well as the 3q29 microdeletion/duplication region. In these cases, inversion formation may serve to create either pre-mutation or protective haplotypes for CNV formation. Depending on the flanking structures, this may “switch” the critical region from predisposing to recurrent inversion, to predisposing to recurrent morbid copy number variation, as previously shown for the 17q2.31 inversion associated with KdV where the toggling events happened during primate evolution (Zody et al. 2008; Itsara et al. 2012). Notably, in the case of the 3q29 inversion our data are consistent with the observation, reported in a current preprint, of 3q29 microdeletions forming on patient haplotypes in direct, rather than the inverted, genomic orientation (Yilmaz et al. 2021). At the genome-wide level, we observe reorganization of >1,000 SD pairs mediated by more than 100 polymorphic inversions—often, but not always, in a complex manner. These observations suggest a vast potential of inversions to prime or protect against subsequent morbid CNV formation, thereby shaping the human landscape of repeat-mediated mutation that is yet to be fully explored.

When combining our data across the autosomes and sex chromosomes, 70% (28/40) of recurrent inversions exhibit ≥10 kbp flanking inverted repeat sequences with high (≥79%) sequence identity (**Table 1**), consistent with a predominant role of NAHR in inversion toggling. On the other hand, for 12% of the fully sequence-assembled balanced inversions we find additional, occasionally complex rearrangements in conjunction with the inversion, likely as a consequence of error-prone DNA repair (Sudmant et al. 2015; Carvalho and Lupski 2016; Collins et al. 2020). Future technological advances allowing to sequence-resolve inversion breakpoints within large SDs will clarify whether error-prone DNA repair contributes to the massive alterations in SD architectures seen at the flanks of recurrent inversions.

### Future considerations and remaining limitations of our work

Integrated haplotype-based assembly of inversions is a prerequisite for inferring mutational recurrence along the human genome. We chose to be conservative in inferring inversion toggling by requiring confirmation of two independent approaches to be certain of recurrent inversion status, with the exception of the Y chromosome that does not recombine meiotically outside of the pseudoautosomal regions. This strategy, thus, may underestimate inversion recurrence, especially if the inverted (or normally oriented) allelic state is subject to some form of selection as is the case for the 17q21.31 inversion polymorphism (Steinberg et al. 2012). Also, we currently base our analysis on 82 unrelated haplotypes; as more genomes are analyzed, inversions confidently classified as a single-origin event in our study may be reclassified as recurrent. An important consideration is that most recurrent inversions are flanked by large inverted SDs (**Table 1**), and that SD-rich genomic architectures are still incompletely resolved by routine long-read sequencing and assembly methods (Ebert et al. 2021). We therefore caution that the sequence assembly of SDs even with phased long-read data can be difficult and additional experiments will be needed to confirm particular SD organizations described in our study. A major challenge going forward will be the complete sequence resolution of large SDs in human genomes. While phased assemblies continue to improve, in our study, Strand-seq was critical for the discovery and genotyping of most large (>100 kbp) inversions (**Fig. 1E**), as well as for resolving inversions embedded in long blocks of duplicated sequence (**Fig. S44**). In addition, orthogonal optical mapping data, such as Bionano Genomics, in concert with the long-read and Strand-seq data, can help to validate more complex structures occurring at the flanks (e.g., chromosome 1p36.13 SDs). Comprehensive characterization of the full spectrum of genomic structural variation in human populations at the sequence level remains an important goal that is likely to be unattainable outside of a multi- platform approach.

## Methods

### Strand-seq data generation and data processing

Strand-seq data were generated as follows. EBV-transformed lymphoblastoid cell lines from the 1KG (1000 Genomes Project Consortium et al. 2015) (Coriell Institute; **Table S1**) were cultured in BrdU (100 uM final concentration; Sigma, B5002) for 18 or 24 hours, and single isolated nuclei (0.1% NP-40 lysis buffer (Sanders et al. 2017)) were sorted into 96-well plates using the BD FACSMelody cell sorter. In each sorted plate, 94 single cells plus one 100-cell positive control and one 0-cell negative control were deposited. Strand-specific DNA sequencing libraries were generated using the previously described Strand-seq protocol (Falconer et al. 2012; Sanders et al. 2017) and automated on the Beckman Coulter Biomek FX P liquid handling robotic system (Sanders et al. 2020). Following 15 rounds of PCR amplification, 288 individually barcoded libraries (amounting to three 96-well plates) were pooled for sequencing on the Illumina NextSeq5000 platform (MID-mode, 75 bp paired-end protocol). The demultiplexed FASTQ files were aligned to the GRCh38 reference assembly (GCA_000001405.15) using BWA aligner (version 0.7.15-0.7.17) for standard library selection. Low-quality libraries were excluded from future analyses if they showed low read counts, uneven coverage, or an excess of ‘background reads’ yielding noisy single-cell data, as previously described (Sanders et al. 2017). Aligned BAM files were used for inversion discovery as described below. On average, there are 68 (median: 57) single-cell Strand-seq libraries per sample (n=44, including newly and previously published Strand-seq data included in the study), with an average 964,021 (median: 820,090) BWA-aligned reads (mapq ≥10) per cell (**Fig. S45**).

### Strand-seq-based inversion discovery

To detect inversions using Strand-seq data, directional composite files were generated for each sample as previously described (Sanders et al. 2016; Chaisson et al. 2019). To automate composite file generation, we implemented a merging protocol based on the breakpointR ‘synchronizeReadDir’ function (Porubsky, Sanders, Taudt, et al. 2020), which locates Watson-Watson (WW) and Crick-Crick (CC) regions in each chromosome and for each cell before building these into the sample-specific composite files. Segmental changes in composite file orientation, suggestive of an inverted allele, were identified using breakpointR (Porubsky, Sanders, Taudt, et al. 2020). To detect both larger and smaller strand-state changes, we used breakpointR in two settings—applying either a window size length of 5 kbp or 20 reads per bin. In both cases, we scaled an initial bin size by multiples of 2, 3, 4, 5, 10 and 20. This resulted in a redundant dataset with putative inversions detected per sample (**Fig. S45**) (mean and median: 144 per sample).

To construct a nonredundant set of Strand-seq inversions, we merged and filtered all detected strand-state changes (putative inversions) in multiple stages as follows: We start out by cropping out each inversion flanks that overlap with highly identical SDs (≥98% identity) or gaps defined in GRCh38. Second, we iteratively merged inversion ranges with ≥50% reciprocal overlap until no more ranges could be merged. Such collapsed ranges were then subjected to a re-genotyping step using the ‘genotypeRegions’ function of the primatR package (Porubsky et al. 2020). Each region in each sample was assigned a genotype: ‘HET’ - approximately equal mixture of plus and minus reads, ‘HOM’ - majority of minus reads, ‘REF’ - majority of plus (reference) reads and ‘lowReads’ - less than 20 reads in a region. Ranges that genotype only as a reference (‘REF’) orientation or have less than 20 (‘lowReads’) reads across all samples were filtered out. Next, we collapsed ranges that share the same genotype across all samples and are embedded with respect to one another. Lastly, for regions that were genotyped only as a ‘HET’ or ‘lowReads’ across all samples, we retained only unique (nonoverlapping) genomic ranges. The same procedure was repeated for window sizes defined by the reads per bin (20 reads per bin as mentioned above), which allowed adding smaller inversions, missed by the larger bin size, to the final Strand-seq inversion callset. Finally, we merged the nonredundant set of inversions created by both window sizes (5 kbp and 20 reads per bin) into a final nonredundant set (n=341) created by the automated procedure described above (**Fig. S46**).

We then manually curated the resultant Strand-seq calls to increase the overall accuracy of the callset, as was done in previous studies (Chaisson et al. 2019; Porubsky et al. 2020). We projected sample-specific composite files onto the UCSC Genome Browser in order to evaluate the mapping of Strand-seq reads inside complex regions of the genome. This procedure led to the addition of 26 inversions and divided 31 regions into more than one inverted event with respect to the automated nonredundant callset (n=341) (**Fig. S47**). This resulted in the final manually curated nonredundant inversion callset (n=419). Among those 419, 39 inversions were marked as false positives mostly caused by a single sample (NA19239) likely due to the extent of background reads in sample-specific composite file (**Fig. S47**). As expected, flagged false positive calls were rather small and genotyped as ‘HET’ (approximately equal mixture of plus and minus reads) as a result of local perturbations in levels of background reads (**Fig. S47**). The manually curated Strand-seq inversion calls were subsequently expanded into the redundant callset (n=6642) and used in the inversion merging process along with PAV- and Bionano-specific inversion callsets (as described below) (**Fig. S46**).

### Inversion discovery from haplotype-resolved long-read assemblies using PAV

Haplotype- phased assemblies (**Data Availability**) were used to generate a long-read-based inversion callset, using the PAV (Ebert et al. 2021) tool, and these assemblies were further utilized to perform sequence-level characterization of inverted sequences. PAV was run on 32/44 samples (64/88 haplotypes) with available phased assemblies. Briefly, PAV aligns each assembled haplotype (2 per sample) with minimap2 (Li 2018) and finds evidence of inversions by analyzing fragmented alignments and aberrant SV patterns created when alignments traverse through an inversion breakpoint. As part of our previous work, variants in assembly collapses were identified using SDA (Vollger et al. 2018) and removed.

### Long-read assembly-based discovery of L1-internal inversions mediated by twin-priming

Non- reference L1 insertion calls previously generated by the HGSVC (Ebert et al. 2021) were subjected to a refined version of the MEIGA-PAV annotation pipeline in order to identify and characterize twin-priming events. First, in order to have all L1 inserts in forward orientation, the reverse complement sequence for every L1 insertion occurring in the minus strand was obtained. Then, poly (A) tails were detected and trimmed for every insert, requiring poly (A) monomers to be at least 10 bp in size, have a minimum purity of 80%, and be located at a maximum distance of 30 bp relative to the insert end. The resulting trimmed inserts were aligned using BWA-MEM 0.7.17- r1188 into a consensus L1 sequence derived from the 632 FL-L1 insertions included in the HGSVC callset. In order to maximize sensitivity for particularly short L1 events a minimum seed length (-k) of 8 bp and a minimum score (-T) of 0 were used. Alignment hits over the L1 consensus were chained based on complementarity in order to identify the minimum set of nonoverlapping alignments that span the maximum percentage of the inserted sequence. A second targeted alignment with BWA-MEM is applied to insert ends that failed to align in the initial alignment round. Based on the alignment chains, L1s are classified as full-length (single hit spanning >99% of the consensus L1), 5’ truncated (single hit spanning ≤99% of the consensus L1), and 5’ inverted (two hits with the first in reverse while the second in forward orientation). Then, the inversion junction conformation for every twin-priming event is determined based on the alignment position over the consensus for the inverted and non-inverted L1 pieces. Blunt joints are characterized by perfect complementary alignments, while overlapping and discontinuous alignments define duplications and deletions at joints, respectively.

Reference L1s were processed similarly as non-reference with two additional preprocessing steps prior to annotation with MEIGA-PAV. RepeatMasker annotations for the GRCh38 genome build were downloaded from the UCSC Table Browser (Karolchik et al. 2004). We noticed that the existence of 5’ inversions frequently led to fragmented annotations, with the inverted and non-inverted sequences being annotated as independent L1s. To correct for this, we merged pairs of L1 annotations adjacent to each other, in opposite orientation and that are complementary at the sequence level. Then, reference L1s were intersected with deletion calls previously generated by the HGSVC (Ebert et al. 2021) in order to select polymorphic L1s that were deleted in at least one of the 64 haplotypes. After reverse complement insertions in the minus strand and trim poly (A) tails, L1 inserts were annotated with MEIGA-PAV.

We successfully inferred the configuration for 93% (1,271/1,362) of L1 polymorphisms, finding that 26% (330/1,271) of them show characteristic 5’ inverted sequences, whereas the remaining are either full-length (405) or 5’ truncated (536). For 7% of L1 polymorphisms (n=91) there was uncertainty regarding the insertion configuration, and these elements were therefore not analysed.

### L1 annotation evaluation via simulations

We generated a simulated dataset, including 9,000 synthetic L1 inserts (**Fig. S48**), evenly distributed among the three possible insertion configurations: full-length (FL), 5’ deleted, or inverted L1. All inserts derived from the same consensus L1 sequence were used as reference for L1 annotation. While the complete consensus was included for FL-L1, random breakpoint positions were sampled for the generation of truncated and inverted events. A single breakpoint located between 10 and 6,013 positions was sampled for 5’ deletions, ensuring a minimum deletion and insertion size of 10 bp. To simulate 5’ inversions, the inversion junction structure was randomly selected among three possible configurations: blunt, deletion, and duplication. Duplication and deletion sizes at the junctions were sampled between 1 and a maximum length of 100 bp and 500 bp, respectively. Similarly, the inversion size was determined based on a random distribution using a minimum inversion length of 10 bp. Then, a random 3’ breakpoint position compatible with the inversion length and junction structure was sampled and the position of the 5’ breakpoint was determined relative to the 3’ breakpoint while taking into account insertion features.

The 9,000 simulated L1 inserts were annotated using MEIGA-PAV and annotations were systematically evaluated using the simulated insertion features as a reference. The predicted insertion configurations were highly consistent with expectations (**Fig. S48B**), with only 18 misannotated insertions, which correspond to insertions with short 5’ deletions misclassified as full-length. Junction conformations were also accurately ascertained (**Fig. S48C**), with 98% (1,016/1,034) duplications, 97% (974/1,008) deletions, and 91% (870/958) blunt joints being concordant. Predicted lengths for inversions, duplications, and deletions at inversion joints were strongly correlated with the expected sizes (**Fig. S48 D-F**). Finally, 76% (4,483/5,940) of all inversion breakpoints were accurately detected, with inaccurate breakpoints having a median deviation of 1 bp (max = 22 bp) (**Fig. S48G**).

### Bionano Genomics–based inversion discovery

We analyzed Bionano Genomics Optical Mapping data by using Saphyr 2^nd^ generation instruments (Part # 60325) and Instrument Control Software (ICS) version 4.9.19316.1. *De novo* assemblies of each sample were obtained using the Bionano Solve v3.5 De Novo Assembly pipeline with haplotype-aware arguments (optArguments_haplotype_DLE1_saphyr_human_downSampleLongestMole.xml) as described previously (Ebert et al. 2021). Using the Overlap-Layout-Consensus paradigm, pairwise comparisons of DNA molecules at least 250 kbp in length, contributing to a coverage of 250X, were generated to create a layout overlap graph and produce initial consensus genome maps. By realigning molecules to the genome maps (alignment confidence cutoff of Bionano p-value < 1e-12) (Anantharaman, Mysore, and Mishra 2004) and by using only the best matching molecules, a refinement step was applied to label positions on the genome maps and to remove chimeric joins. Next, during an extension step, molecules were aligned to genome maps (Bionano p-value < 1e- 12), and the maps were extended based on the molecules aligning past the map ends. Overlapping genome maps were then merged (Bionano p-value < 1e-16). These extension and merge steps were repeated five times before a final refinement was applied to “finish” all genome maps. To identify all alleles, clusters of molecules that were aligned to genome maps with unaligned ends >30 kbp in the extension step were re-assembled to identify potential alternate alleles. To identify alternate alleles with smaller size differences from the assembled allele, clusters of molecules that aligned to genome maps with internal alignment gaps of size <50 kbp were identified, in which case, the genome maps were converted into two haplotype maps. Inversions were identified using the Bionano Solve v3.5 De Novo Assembly pipeline, in which the final genome maps were aligned (Bionano p-value <1e-12) to GRCh38. Manual curation of inversions was performed using Bionano Access (v1.5.2). Optical maps of samples were visually evaluated for inversions not automatically detected by the pipeline. Molecule support for each inversion was evaluated by using the molecule data of each contig containing the inversion. Inversions without molecule support (molecules that span either the entire inversion or anchored to unique labels in proximal and distal inversion breakpoints) were excluded. In addition, inversions identified in centromeric regions and in tandem repeat regions without distinct labelling pattern (direct and inverted configurations of a region of interest show the same labelling pattern) were excluded.

### Bionano Genomics analysis of 1p36 complex region

We analyzed the 1p36.13 region by visual inspection of labelling patterns from optical maps. Segment copies were additionally analyzed in the phased assemblies using BLASTN (version 2.9.0+). Bionano Genomics optical maps were manually evaluated to determine the haplotype in each sample. Single molecules were evaluated using Bionano Access (v1.5.2) to determine whether molecules containing SVs (inversions and CNVs) (n≥3) were anchored to proximal and distal unique regions. Samples with hifiasm (v13) assemblies (n=11) were used as orthogonal support to confirm the haplotypes identified by optical mapping data. The concordance between optical mapping data and sequencing data is estimated to be 89.6% (**Supplementary Methods**).

### Inversion merging into a provisional integrated callset

To create a final nonredundant inversion callset outside of L1 insertions, we merged inversion calls based on different technologies using a previously described procedure (Ebert et al. 2021). Merging of overlapping inversion calls was done in the following priority order: phased assembly-based (PAV), Strand-seq, and Bionano manual callsets. This means that the PAV range is taken first in case two or more inversion calls overlap. In addition, to prevent removing manually curated Strand-seq calls from more complex regions, such as centromeres, we switched off any filtering (applied in (Ebert et al. 2021)) during the merging step. This merging procedure resulted in a provisional inversion callset with 613 genomic regions.

However, in this procedure, a small number of inversion calls made manually using Strand-seq in complex regions of the genome may have been lost, because a PAV inversion call based on an incomplete assembly takes precedence, and thus may lead to a loss of a valid Strand-seq based call. We thus recovered Strand-seq manual calls with less than 50% reciprocal overlap with the merged callset. By doing so, we ended up adding two simple inversions (chr6-26738711-INV- 24388; chr10-79542902-INV-674513) and three inverted duplications (chr16-55798460-invDup- 32830; chr17-19240629-invDup-2318213; chrX-141585258-invDup-102910) to the final provisional merged callset (n=618) (**Fig. S46**). We subsequently continued with the re-genotyping of all regions using ArbiGent as described below. Note that for any PAV call we set to genotype inner breakpoints reported by PAV whenever inner breakpoints were completely embedded within outer breakpoints.

### Inversion genotyping and phasing with ArbiGent

We devised a Strand-seq-based inversion genotyping method, termed ArbiGent, which we employed for three purposes: 1) to unify inversion calls across samples, 2) to verify inversion calls made with other platforms, and 3) to integrate information about inversion loci across samples and accordingly improve the individual callsets. ArbiGent determines inversion genotype likelihoods for genomic loci containing at least 500 bp of sequence uniquely mappable with 75 bp paired-end reads (**Fig. S49**), using strand-specific reads as an input. ArbiGent utilizes an adapted statistical framework previously used for subclonal SV calling in cancer (Sanders et al. 2020). We extended this framework to allow estimating SV genotype likelihoods for DNA segments of choice using Strand-seq data. Based on a Bayesian probability framework that models strand- and haplotype-specific read counts using negative Binomial distributions (Sanders et al. 2020), ArbiGent computes inversion genotype likelihoods for inversions and copy number changes. SV genotype likelihoods derived from individual cells from the same sample are concatenated by summing up log-likelihoods across cells, to result in a combined genotype likelihood estimate per sample and genomic locus of interest. We consider genotype calls made by ArbiGent as ‘high confidence’ if they display a likelihood ratio over a reference state of >10^3^. Integrating genotype labels across samples, ArbiGent additionally assigns labels at the locus level, including ‘potentially FalsePositive (FP)’ if the locus is never seen inverted in any sample, or ‘alwaysComplex’ if the locus was not seen in a non-complex (or reference) state in any sample. In GRCh38, 489 out of 615 (79.5%) inversions in the initial discovery set were assigned a ‘passing’ label (‘pass’,‘noreads’,‘lowconf’,‘INVDUP’,‘MISO’), with the ratio increasing for inversion loci called by at least two independent techniques (75 out of 77, 97.4%) (**Fig. S50**).

ArbiGent supports haplotype phasing of events based on the StrandPhaseR method, which infers phase information from Strand-seq reads (Porubský et al. 2016; Porubsky et al. 2017). We additionally synchronized the haplotype assignments per chromosome (H1 vs. H2) post-hoc to the long-read-based phased genome assemblies (in the subset of samples [35/44] where such assemblies were available). This is done by comparing heterozygous SNP sites called by StrandPhaseR (Porubsky et al. 2017) to those called by PAV, and reversing the ArbiGent phase assignment for all chromosomes where the PAV and Strand-seq SNPs are phased orthogonally. This procedure was applied to all inversions on 796/805 chromosomes (35 independent samples × 23 chromosome sets), while the remaining nine displayed potential flaws in PAV phasing in which case ArbiGent calls were reported using their original phase (**Fig. S51**). Using this procedure, we genotyped all 618 inversions, of which 615 were used for subsequent analysis (three inversions from unassigned fragments of the GRCh38 reference were dropped: chr14_GL000225v1_random- 107057-INV-5515, chr17_GL000205v2_random-160765-INV-1685, chrUn_KI270743v1-150894-INV-11414; **Fig. S46**).

### Inversion filtering and generation of the final integrated callset

We applied a number of filters to remove low-quality inversion calls and generate the final integrated inversion callset. First, we removed calls that were genotyped by ArbiGent as ‘alwayscomplex’, ‘alwayscomplex-INVDUP’ or ‘FP’ (false positive). Second, we removed any remaining inversions unique to the Strand-seq callset that were flagged as either ‘FP’ (false positive) or SD (‘segmental duplication’). Third, we removed any unique automated Bionano call (reported by automated Bionano procedure) with less than 90% reciprocal overlap with the manually curated Bionano inversion callset (**Methods**). This was done due to the imperfect overlap between the automated and manually curated Bionano callsets, and in order to prevent introducing false positive inversion calls (**Fig. S52**). Lastly, we dropped inversion calls with 90% or more reciprocal overlap with another inversion call within the callset, which brought the number of inversions to 418 (**Fig. S52**). We marked putative misorients as regions defined as ‘miso’ by ArbiGent or marked as putative misorient during the manual curation of Strand-seq callset (all carriers show homozygous inversions). Any call that showed at least one clear heterozygous genotype was not marked as putative misorients.

To finalize the callset we manually evaluated dot plot alignments for all 418 inversions, using the phased assemblies (**Supplementary Methods**). We identified contigs spanning both breakpoints for 183/418 (44%), predominated by smaller inversions below 200 kbp (**Fig. S53**). We used the alignments to verify inversion status and to annotate repeats at the breakpoints as well as other SVs at the flanks, such as insertions, deletions, and duplications. We used a 50 bp lower cutoff in reporting homology and SVs. Finally, we intersected all homologous repeats with mobile elements present in RepeatMasker v4.1.2 ((Tarailo-Graovac and Chen 2009). Using this approach, we identified inversions (n=31) likely driven by mobile elements (both inversely oriented homologous repeats displayed >80% reciprocal overlap with an individual mobile element of the same class). Using the phased assemblies, we then adjusted inversion breakpoint positions. After breakpoint adjustment, we re-genotyped all inverted regions once more using ArbiGent. We then manually checked inversions for potentially redundant calls, which were removed from the callset. This led to the removal of six redundant calls (variant IDs: chr6-167197698-INV-159879, chr8-2235761- INV-247546, chrX-52472287-INV-68115, chrX-155384040-INV-73061, chr10-46986413-INV-

193930, chr10-79526936-INV-290309). We also removed calls flagged as false positives during the manual dot plot evaluation and with no support by Strand-seq data (n=13), resulting in the final callset of 399 inversions. Finally, to avoid reporting low-confidence genotypes for small inversions discovered only by PAV, we replaced the ArbiGent genotypes with PAV genotypes for these small events if the ArbiGent genotype was labelled as “low confidence”.

We assigned each inversion call into the following classes: ‘inv’ (balanced), ‘invDup’ (inverted duplication), ‘miso’ (putative misorientation), or ‘complex/low confidence’ (complex) based on ArbiGent genotypes followed by manual curation. Sites with at least one high-confidence inverted allele are marked as balanced inversions (‘inv’), misorients (‘miso’) were marked as described above, inverted duplications (‘invDup’) were marked based on ArbiGent prediction, and the rest of the calls fall into the complex or low-confidence category (‘complex/low confidence’). We emphasize that such extensive manual curation of our callset is crucial to deliver accurate and confident inversion callset because many inversions lie within the most complex repeat-rich regions of the human genome.

### Phasing and correction of chromosome-length inversion haplotypes

While Strand-seq is by design well suited to detect inversions and can be used for long-range (chromosome-length) haplotyping, the haplotypes constructed by StrandPhaseR (Porubský et al. 2016; Porubsky et al. 2017) over the inverted region are not in the correct phase with the rest of the chromosome. This is because inversions change the directionality status (plus or minus) of Strand-seq reads with respect to the surrounding regions. The scope of the problem differs for homozygous and heterozygous inversions. Homozygous inversions appear as a complete switch of haplotypes in comparison to the haplotypes from uninverted regions. This is caused by a switch in directionality of Crick (plus) and Watson (minus) reads with respect to the uninverted regions. Such a switch in haplotypes inside a homozygous inversion is easily corrected by flipping such haplotypes. In contrast, heterozygous inversions are more difficult to correct as only one strand is inverted, and thus, either the Watson or Crick strand changes its directionality over the inverted region (**Fig. S2**). This creates a mixing of alleles in heterozygous state so the heterozygous inversions need to be phased *de novo* using Strand-seq cells that inherited either only Watson or Crick strands from each parent for a given chromosome. In such cells, heterozygous inversions appear as an equal mixture of Crick and Watson reads as only one strand (one haplotype) is inverted. Such cells are informative and can be used for unambiguous phasing for a given inverted region, since the Crick and Watson reads are coming from different parental homologs. Inverted haplotypes were assigned to a respective parental homolog based on the phasing of reads from Strand-seq cells that inherited either Watson or Crick strands from each parent. We implemented the functionalities for correcting the phase of inverted sequenced in the R package StrandPhaseR (v0.99), in a new function called ‘correctInvertedRegionPhasing’. We supplied this function with the sample-specific inverted regions reported in this study, in order to correct each of them using the given function parameters (recall.phased = TRUE, het.genotype = ’lenient’, pairedEndReads = TRUE, min.mapq = 10, background = 0.1, lookup.bp = 1000000, bsGenome = BSgenome.Hsapiens.UCSC.hg38, assume.biallelic = TRUE).

After inversion phase correction, chromosome-length haplotypes based on Strand-seq data were used to guide phasing of the long-read assemblies, by executing a previously described integrative phasing framework (Porubsky et al. 2017). Integrative phasing was completed using WhatsHap (version 0.18) and subsequently linkage disequilibrium was calculated using PLINK (version 1.9), with a window size of 200 kbp. For integrative phasing, we used a defined set of variant positions (available at http://ftp.1000genomes.ebi.ac.uk/vol1/ftp/data_collections/1000G_2504_high_coverage/working/20201028_3202_phased/). Finally, we re-genotyped the sample VCFs based on the long-read BAM files.

### Inversion site verification using ONT reads

We used publicly available ONT reads from three samples—HG002, HG00733 and NA19240 (**Data availability**)—in order to verify inverted regions in our integrated inversion callset outside of L1 insertion sequences (n=399). First, we mapped ONT reads onto the reference genome (GRCh38) using minimap2 (Li 2016) (version 2.20-r1061) using the following parameters: --secondary=no -z 400,0 -r 100,1k. All alignments were reported in the PAF format. We processed each alignment that was no further than 1 kbp from the predicted inversion breakpoints. For further analysis we kept only those ONT reads that showed a split alignment with at least one plus and one minus oriented alignment. Next, we calculated the fraction of bases contributed by inverted and direct alignments inside and outside of the inversion range. We considered an inversion to be supported by ONT reads if there were at least three split-read mappings. We also required that the fraction of inverted base pairs inside an inverted region is at least 0.5 higher than the fraction of inverted base pairs outside of the inverted region (**Fig. S54**).

### PCR validation

As a further line of validation for the callset, we subjected 10 randomly selected breakpoint-resolved inversions (length: 0.8–366 kbp) to site and genotype validation via PCR. Primers were designed using a computational SV validation primer design pipeline, as previously described (Sudmant et al. 2015), which is in turn based on the Primer3 method (Koressaar and Remm 2007) and available at https://github.com/zichner/primerDesign. Briefly, the pipeline uses an iterative approach to extract two uniquely mapping, inversely oriented primers pairs per inversion, with each pair mapping to opposite sides of one inversion breakpoint. Utilizing sequences generated with primerDesign, combinations of primer pairs were then systematically tested for PCR amplification in supposed inversion carriers and non-carriers to validate genotypes. We designed four primers per inversion: primer 1 (P1) and primer 3 (P3) are designed flanking the leftmost breakpoint of the inversion site and would amplify a region spanning the breakpoint in a reference (direct) orientation. Primer 2 (P2) and primer 4 (P4) are analogously designed flanking the rightmost breakpoint and would amplify a region spanning the breakpoint in a reference orientation. In an inverted locus, P2 and P3 would switch orientation and the combinations P1+P3 or P2+P4 would no longer yield a PCR product, as both primers would be in the same orientation. In an inverted orientation, P1+P2 and P3+P4 combinations would be productive and yield an amplification around the breakpoint. PCR primers were obtained from Sigma-Aldrich at 100 µM concentration in H2O. Genomic DNA was either ordered from Coriell or extracted from cultured cell lines with Qiagen’s QIAamp DNA Blood Mini Kit and set to 10 ng/µl concentration. PCR was done with ThermoFisher’s Phusion Human Specimen Kit using the 20 µl reaction volume consisting of 10 µl of 2x reaction buffer, 0.4 µl Phusion polymerase, 7.6 µl of H2O, 1 µl of 10 µM primers (final concentration 1 µM), and 1 µl of 10 ng/µl genomic DNA. Cycling conditions were: 98C 5min, 34x cycles of 98C 1sec, 63C 5sec, 72C 2min, 72C 5min, 4C hold. Amplified fragments were visualized in 2% agarose gels containing SybrSafe (ThermoFisher).

### Identifying potential inversion carriers using PanGenie

In order to find other potential carriers of the pericentromeric inversion on chromosome 2 detected in NA19650 and the 5 Mbp inversion on chromosome 15 detected in HG02492, we considered SNP alleles present in the inversion haplotypes of these two samples as follows. We used the HGSVC freeze 4 SNP genotypes (Ebert et al. 2021) available for all 3,202 1KG samples (Byrska-Bishop et al. 2021) (http://ftp.1000genomes.ebi.ac.uk/vol1/ftp/data_collections/HGSVC2/release/v2.0/PanGenie_res ults/) to determine which of these SNPs are rare (allele frequency ≤0.01 across unrelated samples). In HG02492 we identified 103, and for NA19650 we detected 333 such rare SNPs within the respective inverted segments. We then counted these rare alleles in the genotypes of all 3,202 samples. Those samples that share a high number of rare SNPs with the respective inversion haplotype were considered potential carriers of the inversion. For the inversion on chromosome 15, we identified four samples sharing a high number of rare SNP alleles with HG002492: HG002491 (102/103 alleles in common; 99%; mother of HG02492), HG02784 (101/103; 98%), HG02725 (74/103; 72%), and HG03639 (74/103; 72%). For the pericentromeric inversion on chromosome 2, sample NA19648 (mother of NA19650) shared 330/333 (99%) rare alleles with the respective haplotype-resolved segment in NA19650.

### FISH validation

Metaphases were obtained from eight human lymphoblast cell lines (NA19648, NA19650A, HG03639, HG02784, HG02725, HG02491, HG02492 and NA12878 as control).

Two-color FISH experiments were performed using human fosmid (n=2) or BAC (n=2) clones directly labelled by nick-translation with Cy3-dUTP (PerkinElmer) and fluorescein-dUTP (Enzo) as previously described (Lichter et al. 1990), with minor modifications. Briefly, 300 ng of labelled probe was used for the FISH experiments; hybridization was performed at 37°C in 2xSSC, 50% (v/v) formamide, 10% (w/v) dextran sulphate, and 3 mg sonicated salmon sperm DNA in a volume of 10 mL. Post-hybridization washing was at 60°C in 0.1xSSC (three times, high stringency). Metaphases were simultaneously DAPI stained. Digital images were obtained using a Leica DMRXA2 epifluorescence microscope equipped with a cooled CCD camera (Princeton Instruments). DAPI, Cy3, Cy5, and fluorescein fluorescence signals, detected with specific filters, were recorded separately as grayscale images. Pseudocoloring and merging of images were performed using Adobe Photoshop software. Since the tested inversions were >2 Mbp, two-color FISH on metaphase chromosomes was performed using two probes within the inverted region and the centromere as anchor.

### Excess in common inversion polymorphisms in the genome

Since we were evaluating inversions excluding misorients (found in all haplotypes), we also excluded variants found in all haplotypes from the SV and single-nucleotide variant (SNV) callsets to avoid biasing allele frequency and growth rate comparisons. Singletons are defined as variant calls with an allele count of 1. Statistics were computed per haplotype (two haplotypes per sample). Callset growth rate was computed for each haplotype by taking the number of singletons and dividing by the callset size less the number of singletons, which is the proportion of growth if the whole callset was constructed and that haplotype added. This was computed for each haplotype, and the mean growth rate was reported. The supporting p-value was computed over the growth rate for all samples, computed for each variant type, and testing two variant types with a two-tailed Student’s T-test assuming independence. By comparison, we computed 0.48% callset growth for SV insertions and deletions, which did not differ significantly from the SNP growth rate (*P=*0.45, t-test).

We noticed an excess in common (minor allele frequency [MAF]>5%) inversion alles (67%) compared to other SV classes (48%) and SNPs (47%), which likely explains the callset growth rate. A test of significance of allele frequency cutoffs was conducted by splitting the callset by allele frequency (<0.05 and ≥0.05) and comparing the counts with a two-tailed Fisher’s exact test. For this test, we corrected the SV insertion and deletion (SV INS/DEL) callset variant sizes from Ebert et al. to eliminate differing distributions from smaller SVs, and we eliminated all shared SV INS/DEL variants to maintain compatibility with the inversion callset. Before correction, 52% of the SV INS/DEL calls (n: INV=292, INS/DEL=105,913) had an allele frequency ≥5% with a p-value of 7.85e-08, (SV INV vs SV INS/DEL split at 5% allele frequency, two-tailed Fisher’s exact test). When SV INS/DEL variants were restricted to events 300 bp or greater (to better match the callsets by size) as the minimum inversion size, (n: INV=292, INS/DEL=38,734), we observed 48% with allele frequency ≥5% *(P*= 2.63e-11), which become 29% (*P*= 1.83e-08) for SV INS/DEL ≥10 kbp (n: INV=203, INS/DEL=906). This strongly supports the observation that uncommon inversion alleles (<5% AF) are significantly depleted when compared to SV insertions and deletions of similar size and that the results remain significant whether or not they are corrected for similar SV size. This suggests an excess of common inversion polymorphisms in the human genome.

### SNP-based analysis for inversion recurrence

In this study, we developed a statistical tiSNPs- based approach that detects, using the haplotype-resolved Strand-seq data, evidence of inversion recurrence by individually considering the occurrence of biallelic SNPs within an inverted locus (**Fig. 3B**). The decision of whether a SNP suggests inversion recurrence is made based on how often each of its alleles occur in an inverted/non-inverted haplotype across all samples. On the basis of aggregated evidence across all SNPs, the inversion is then termed as ‘recurrent’ or ‘single-event’. We based this analysis on the set of 279 ‘balanced’ inversions from the autosomes and chromosome X.

The analysis steps are as follows:

1. For each biallelic SNP within an inversion, the number of Strand-seq reads in Watson (W) and Crick (C) orientation were recorded. The reads were further filtered for quality by removing secondary alignments, duplicates, and reads with mapping quality lesser than ‘10’. The read counts were maintained individually for each single cell per sample. For this analysis, we only consider biallelic SNPs with allele frequency ≥5% because with rarer SNPs the method does not have sufficient power to detect evidence of recurrence. This SNP filtering led to the removal of 27 inversions, leaving us with 252 that could be further tested.
2. Using the background/normal cell state, these strand notations were translated to ‘forward/non-inverted’ and ‘reverse/inverted’ notation. For example, if the background cell state is ‘CC’, all the Watson reads mapping to the SNP would be termed as ‘inverted’ while all the Crick reads would be termed as ‘non-inverted’. For ease of comparison to the normal cell state, only ‘WW’ and ‘CC’ cells from each sample were considered for further analysis.
3. At this point, we had a record for each ‘within inversion’ SNP per single cell, indicating how often we observed each SNP allele in the ‘inverted’ and ‘non-inverted’ state. These occurrence counts were then aggregated first across all single cells and then across all samples, resulting in a table that stored for each SNP the occurrence of each possible SNP-inversion haplotype configuration.
4. The next step was to identify tiSNPs. Theoretically, observing each SNP allele in both ‘inverted’ and ‘non-inverted’ haplotype at least once indicates that the inversion recurred, but to account for the ‘background’ reads in the wrong orientation (∼5% in Strand-seq data), a SNP was termed as a ‘toggling-indicating SNP’ or tiSNP, only if each of the four possible SNP-inversion configurations was seen at least thrice, i.e., each of its alleles had at least ‘3’ ‘inverted’ and ‘non-inverted’ reads mapped to them.
5. For a quantitative assessment, for each inversion, a record of the fraction of tiSNPs compared to the total number of considered SNPs was maintained.

For 49/252 inversions, we observed at least one tiSNP. To make sure our approach is detecting true evidence of recurrence, the analysis was applied to a control set consisting of random, non-inverted regions of the human genome with the same size distribution as the inversion callset. Only 0.02% of the randomized intervals showed evidence of recurrence with the fraction of tiSNPs being extremely low (≤0.002). Using this approach, there is an extremely low possibility of seeing a recurrence signal randomly, which supports the claim that it enables inferring ‘true’ signals of inversion recurrence.

*Influence of flanking inverted repeats on inversion recurrence:* An important feature distinguishing recurrent and nonrecurrent inversions is their flanking sequence. Mechanistically, inversions are usually mediated by homologous inverted sequences flanking both inversion breakpoints. Additionally, it is hypothesized that the longer the stretch of flanking inverted repeat, the higher the rate of inversion recurrence. To test whether our analysis supports this hypothesis, inversions labelled as ‘recurrent’ were compared to the ones labelled as ‘single-event’, in terms of length of the longest flanking inverted repeat sequence. Only the repeats with one end lying within 20 kbp (-10 kbp to +10 kbp around each annotated breakpoint) and extending up to 70 kbp flanking region were considered. The inversions showing evidence of recurrence turned out to be clearly enriched for longer flanking inverted repeats with fraction of tiSNPs increasing with increasing length, while the ones where we did not observe any recurrence signal showed enrichment for shorter flanking inverted repeats (**Fig. S55**).

*Influence of inversion length on inversion recurrence:* Another quality check was to make sure that the analysis is not being driven by inversion length, because, keeping in view the approach formulation, it is more likely to observe a recurrence signal in longer inversions with more SNPs as compared to small inversions with fewer SNPs. Additionally, the length of flanking inverted repeats is believed to be directly associated with inversion length. Therefore, to add statistical support to the claim that the observed relationship between length of flanking inverted repeat and the fraction of tiSNPs does not involve inversion length as a confounder, a multiple linear regression model was used. The model expressed the fraction of tiSNPs as a function of inversion length, length of the longest flanking inverted repeat and major allele frequency. Major allele frequency was considered here, because for inversions where either of the alleles is rare, the statistical power to detect recurrence is reduced as compared to inversions with balanced allele frequency. The regression results clearly suggested that the recurrence signal detected by the analysis is primarily influenced by the length of flanking inverted repeats and major allele frequency (*P=* 7.5e-04 and *P*= 2.65e-06, respectively), with inversion length having no significant influence (*P*= 8.35e-01).

### Haplotype-based coalescent approach for detecting inversion recurrence

As a second approach, we developed a haplotype-based coalescent approach that considers four lines of empirical evidence for inversion recurrence, using phylogenetics and population genetics methods: 1) haplotype-based PCA, 2) haplotype identity by state, 3) window-based phylogenetic tree reconstruction, followed by bootstrap analysis, and 4) reconstruction of ancestral recombination graphs using Relate (Speidel et al. 2019). Input VCFs were created using the procedure described in Methods section ‘Phasing and correction of chromosome-length inversion haplotypes.’ Individuals who we failed to unambiguously assign an inverted haplotype were removed from any analysis. In addition, as a QC filter we excluded SNVs mapping to SDs (sequence identity >98%) and heterochromatic satellites including centromeres and telomeres. To ensure the quality of our analysis, we focused on 127 inversions having sufficient unique sequence and at least 10 SNVs within an inversion to construct haplotypes. We describe each method separately and provide a detailed description of the procedures.

- **PCA:** We performed a haplotype-based PCA following the description of Browning et al. (Browning et al. 2016). Briefly, each haplotype base was encoded numerically, where 0 and 1 (and 2 if needed) represent ancestral (using the chimpanzee assembly as the outgroup (Kronenberg et al. 2018)) and derived alleles, respectively. We used the R package, irlba (v2.3.3), to calculate and visualize principal components and keep track of direct and inverted alleles.
- **Haplotype identity by state:** Identity by state is defined as the proportion of matches between a certain pair of haplotypes. Pairwise differences between haplotypes were visualized and grouped based on inverted versus direct orientations.
- **Ancestral recombination graph reconstruction:** We used the software Relate (Speidel et al. 2019) to reconstruct the local genealogy of an inversion locus. The method infers underlying coalescent events consistent with the observed data under the infinite-sites model, taking into account recombination, and thus, recapitulates the multi-locus genealogy of the genomic region. We used a mutation rate of 1.25 × 10^!%^per base per generation and set the haploid effective population sizes as 10,000 (Gutenkunst et al. 2009) and 6,900 (Veeramah et al. 2014) for autosomal and X chromosome loci, respectively. We projected the inversion genotype onto the resulting trees and inferred the number of inversion events using Fitch’s algorithm for homoplasy (Fitch 1971). The rate of inversion at each inversion locus is defined as the estimated number of inversion events divided by the total tree length in generations. Note that for a given tree, a monophyletic inversion group indicates a single origin of these inverted haplotypes, while a polyphyletic group of inverted haplotypes suggests that these inverted haplotypes are derived from more than one common ancestor and, thus, recurrent. To evaluate the consistency of topology across inferred trees, we used generalized Robinson-Foulds distances implemented in the R package TreeDist (v2.1.1).
- **Phylogenetic tree reconstruction:** For phylogenetic tree reconstruction, we applied the maximum likelihood-based software IQ-TREE (v2.1.3) using the following options: ‘- keep-ident -redo -bb 1000 -m MFPMERGE --date *date.file* --date-options “-u 0” --clock- sd 0.4 --date-tip 0 ‘. We assumed six million years of divergence between human and chimpanzee (specified in the date.file). Firstly, for each inversion locus, we inferred phylogenetic trees for the entire locus and for 100 block-bootstraps and estimated the number of inversion events for individual trees to assess confidence. Secondly, we sliced the region into 2,000 and 20,000 bp windows for loci smaller and greater than 20,000 bp, respectively, and inferred a local phylogenetic tree for each window. We again used the Robinson-Foulds distances to evaluate the consistency of tree topology among inferred trees across the locus and computed the number of independent inversion events for each local tree. For each tree, the number of events was divided by the total tree length to obtain inversion rates.
- Finally, for each inversion under consideration, we computed three intervals to measure the uncertainty of the inferred inversion rates using 1) a 95% central interval, computed based on the 2.5 and 97.5 percentiles of the estimates from all marginal trees inferred by Relate at a locus; 2) a similar 95% central interval computed from the local trees built using the window-based phylogenetic tree reconstruction; and 3) a 95% confidence interval constructed using the 100 block-bootstrap trees for the entire inversion locus. We determined that an inversion is recurrent if all three intervals indicate at least two independent origins of the inversion. Unless mentioned otherwise, for each inversion tested, we reported the 95% central interval computed based on the inference of Relate throughout the paper as this interval tends to be wider and therefore more conservative.

### Chromosome Y inversion genotyping

ArbiGent, at present, is not well tailored to genotype haploid chromosomes such as chromosome Y in males. Because of this caveat we decided to re-genotype reported chromosome Y inversions (n=15) using Strand-seq data only based on the binomial distribution of Crick (plus) and Watson (minus) reads. For this purpose we used the R function ‘genotypeRegions’ implemented in R package primatR (Porubsky et al. 2020). We required a minimum of five Strand-seq reads (min.reads = 5), in order to report a genotype and allowed for 10% of background reads (alpha = 0.1). Genotypes for chromosome Y inversions are reported as a supplementary table (**Table S9**).

### Construction and dating of Y phylogeny and Y inversion rate estimation

We called the genotypes of 17 samples (16 males included in the current study plus NA19384 used to root the Y phylogenetic tree) jointly from the 1KG Illumina high-coverage data using the ∼10.3 Mbp of chromosome Y sequence previously defined as accessible to short-read sequencing (Poznik et al. 2013). BCFtools (v1.9) was used with minimum base quality and mapping quality 20, defining ploidy as 1, followed by filtering out SNVs within 5 bp of an indel call (SnpGap) and removal of indels. Additionally, we filtered for a minimum read depth of 3. If multiple alleles were supported by reads, then the fraction of reads supporting the called allele should be ≥0.85; otherwise, the genotype was converted to missing data. Sites with ≥6% of missing calls across samples were removed using VCFtools (v0.1.16). After filtering, a total of 10,407,641 sites remained, including 5,494 variant sites.

The Y haplogroups of each sample were predicted as previously described (Hallast et al. 2021) and correspond to the International Society of Genetic Genealogy nomenclature (ISOGG, https://isogg.org, v15.73) (**Table S9**). We used the coalescence-based method implemented in BEAST (v1.10.4 (Drummond and Rambaut 2007) to estimate the ages of internal nodes in the Y phylogeny. A starting maximum likelihood phylogenetic tree for BEAST was constructed with RAxML (v8.2.10 (Stamatakis 2014)) with the GTRGAMMA substitution model using all sites. Markov chain Monte Carlo samples were based on 100 million iterations, logging every 1000 iterations. The first 10% of iterations were discarded as burn-in. A constant-sized coalescent tree prior, the HKY substitution model, accounting for site heterogeneity (gamma) and a strict clock with a substitution rate of 0.76 × 10^−9^ (95% confidence interval: 0.67 × 10^−9^– 0.86 × 10^−9^) single-nucleotide mutations per bp per year was used (Fu et al. 2014). A prior with a normal distribution based on the 95% confidence interval of the substitution rate was applied. A summary tree was produced using TreeAnnotator (v1.10.4) and visualized using the FigTree software.

In order to estimate the inversion rate, we counted the minimum number of inversion events that would explain the observed genotype patterns in the Y phylogeny. A total of 4,419 SNVs called in the set of 16 analyzed males and Y chromosomal substitution rate from above was used. A total of 126.4 years per SNV mutation was then calculated (0.76 × 10^−9^ × 10,407,641 bp)^-1^, which was converted into generations assuming a 30-year generation time (Fenner 2005). Each SNV thus corresponds to 4.21 generations, translating into a total branch length of 18,623 generations for the 16 samples. For a single inversion event in the phylogeny this yields a rate of 5.37 × 10^−5^ (95% CI: 4.73 × 10^−5^ to 6.08 × 10^−5^) mutations per father-to-son Y transmission. The confidence interval of the inversion rate was obtained using the confidence interval of the SNV rate.

### Detection of nested inversions and inversions showing signs of breakpoint reuse

We first determined inversions that are completely embedded within another inverted range in our callset (n=33). In addition to that, we compared genotypes of all possible pairs of inversions and kept those that consistently share the same genotype across all samples (n=17). We compiled these two sets of inversions into a nonredundant candidate list of inversions with potentially shifted breakpoints (n=19) (**Table S10**). We removed any sites that involve putative misorients. We manually inspected binned read counts of Strand-seq data over each candidate region and selected three of the most confident regions where an inversion breakpoint shift is plausible. We also selected one region where we predict a nested inversion event (**Table S10**).

### Inversions changing the orientation of SD pairs

We devised a computational approach that systematically scans all identified inversions for their potential to change the relative orientation of pairs of SDs in the genome, by inverting one SD out of a pair. Starting from an annotated set of 69,906 annotated SDs of >90% sequence identity obtained through the UCSC Table Browser (Karolchik et al. 2004), we selected SDs longer than 10 kbp and with homologous partners on the same chromosome (Mefford and Eichler 2009), resulting in 7,672 SDs, representing 3,795 one-to-one pairs. We then identified SD pairs in which one but not both partners are embedded within the same inversion, yielding 2,265 SDs (1,094 SD pairs). Filtering on the level of inversions next, we identified 79 inversions flipping the orientation of at least one SD pair, with a median of eight pairs reoriented. Out of these, we identified 29 inversions that predominantly (>90% of flipped SD pairs weighted by length) affect SD pairs in direct or in inverse orientation. We classify these inversions as ‘potentially protective’ (n=9) and ‘potential pre-mutational state’ (n=20), respectively. Morbid CNVs from the decipher database (Bragin et al. 2014) were additionally intersected with this set of 29 inversions, identifying 6 inversions overlapping with 8 unique morbid CNVs (**Table S11**).

### RNA-seq data processing and mapping

Before RNA-seq (**Data Availability**) read mapping, adapters and low-quality reads and bases were removed using Trim Galore (Andrews et al. 2015). The remaining reads were mapped to GRCh38 using STAR aligner in 2-pass mode (Dobin et al. 2013). Leveraging sample-specific SNP-calls based on GRCh38 reported previously in Ebert et al. (2021), alignment to this genome reference was performed with WASP filtering (van de Geijn et al. 2015) to mitigate allelic mapping bias. To prepare the data for haplotype-unaware eQTL analysis, reads were quantified with respect to GENCODE v35 genome annotation (Frankish et al. 2019) using featureCounts (Liao, Smyth, and Shi 2014). Read counts were normalized using the TMM normalization implemented in edgeR (Robinson, McCarthy, and Smyth 2010) and transformed to the transcripts-per-million (TPM) metric.

### eQTL mapping

The set of 399 inversions outside of L1 internal sequences was tested for association with expression in nearby genes together with a set of 41,833 deletions, 66,825 insertions, and 16.4 million SNPs as previously reported in Ebert et al. (2021). We first identified expressed genes (TPM > 0.5 in at least 5 samples) located in a window of 2 Mbp centered on each inversion breakpoint. This resulted in 4,469 genes to be tested for association with all variants overlapping a 2 Mbp window centered around the gene. All variants with MAF ≥1% were considered, resulting in a total of 13.4 million gene-variant pairs to test.

eQTL tests were performed using a pipeline based on nonlinear mixed models, implemented in LIMIX (Lippert et al. 2014). PLINK (v.1.90) was used to estimate genetic ‘kinship’-matrices between samples based on SNP and indel variants, which served as a basis for population principal components (PCs) used as latent factors in the model. As additional cofactors, we used PCs of the gene expression matrix to account for remaining systematic expression biases. We found n=4 covariates for bias correction (Expression-PC1, Population-PC1&2, Sex) to maximize the number of discovered eQTLs, with larger numbers of covariates showing signs of overfitting due to the relatively low sample size. The results of the eQTL mappings were initially corrected for multiple testing both on the level of tested variants per gene and the level of number of genes tested, as described in Ebert et al. (2021). Variant-level correction was performed via genotype permutations as implemented in LIMIX, while the number of genes was corrected for using a Storey Q-value– based procedure. Using this rigorous global correction, we find 166 globally significant eQTLs (1 INV, 1 DEL, 164 SNVs). In parallel to this all-variant approach, we also analyze inversion eQTLs separately, allowing us to reduce the number of tests to correct for by a factor of ∼6,600 and recover significant inversion eQTLs that would otherwise be overshadowed by the large number of variants tested. In this approach, we used Benjamini-Hochberg correction on the level of inversion-gene tests, yielding 11 globally significant (FDR < 0.2) inversion eQTLs. Lead eQTLs in all cases were determined by comparing raw p-values of all variants of a given gene.

### Enrichment analysis of inversions intersecting morbid CNV regions

We tested whether various subclasses of inversions outside of L1-internal sequences (n=399), namely balanced inversions (n=292), consensus single-inversion events (n=61), and consensus recurrent inversion events (n=32) are enriched in the vicinity of known pathogenic CNVs (redundant set, n=155) (Cooper et al. 2011; Coe et al. 2014; Bragin et al. 2014). We used the R package regioneR (Gel et al. 2016) with its function ‘permTEST’ to perform permutation testing (n=10,000 permutations). At each permutation, we randomized the position of each inversion in every tested subgroup using regioneR’s function ‘circularRandomizeRegions’. This way the relative distance of each inversion is kept, as inversion occurrence on each chromosome is not completely random and highly depends on the underlying SD architecture. At each permutation, we counted the number of inversions overlapping with the redundant set of morbid CNVs, allowing for a 50 kbp gap between each set of coordinates (to account for the fact that recurrent inversion as well as sites of recurrent microdeletions and microduplications are often flanked by an extensive, hard-to-penetrate SD architecture).

## Acknowledgements

The authors thank Simone Köhler and Tonia Brown for valuable comments on this manuscript as well as editing, and Marc Jan Bonder for providing valuable advice on eQTL mapping. We thank the Genomics Core Facility and the IT Services at the EMBL for valuable technical assistance. Funding for this research project came from the following grants: National Institutes of Health (NIH) U24HG007497 (to C.L., E.E.E., J.O.K., T.M.), U01HG010973 (to T.M., E.E.E., and J.O.K.), as well as R01HG002385 and R01HG010169 (to E.E.E.), the German Federal Ministry for Research and Education (BMBF 031L0184 to J.O.K. and T.M.), and the European Research Council (ERC Consolidator grant 773026 to J.O.K.). Support for the work also came from the European Molecular Biology Laboratory (J.O.K, P.Hasenfeld, E.B.), and the EMBL International PhD Programme (W.H.). E.E.E. is an investigator of the Howard Hughes Medical Institute. B.R.M. is supported by a Bridging Excellence Fellowship provided by the Life Science Alliance. P.Hsieh. is supported by the NIH Pathway to Independence Award (NHGRI, K99HG011041). C.R.B. and P.A.A. are supported by NIH NIGMS R35GM133600 and NCI P30CA034196. F.Y., P.Hallast. and Q.Z. are supported by NIH U24HG007497. Deep Illumina WGS data from the 1KG samples were generated at the New York Genome Center with funds provided by NHGRI Grants 3UM1HG008901-03S1 and 3UM1HG008901-04S2.

## Author contributions

Conceptualization, D.P., A.D.S., T.M., E.E.E., J.O.K.; Methodology, Software, D.P., W.H., H.A., P.Hsieh, B.R.M., M.S.; Formal analysis, D.P., W.H., H.A., P.Hsieh, B.R.M., F.Y., J.E., P.Hallast; Investigation, D.P., W.H., H.A., P.Hsieh., B.R.M, A.D.S., M.S. C.R.B.; Resources, HGSVC, Q.Z., C.L., P.Hasenfeld, A.D.S., T.M., E.E.E., J.O.K.; Computational support, W.T.H., P.A.A., B.H., D.S.G.; Validation, F.A.M.M., P.E., E.B., F.A.; Writing, D.P., W.H., H.A., B.R.M., E.E.E., and J.O.K. with input from all authors.

## Software availability

Main software packages used in this study: primatR (https://github.com/daewoooo/primatR)

breakpointR (https://github.com/daewoooo/breakpointR)

StrandPhaseR (https://github.com/daewoooo/StrandPhaseR, branch=devel)

ArbiGent (https://github.com/friendsofstrandseq/pipeline/tree/arbitrary-segments)

PAV (https://github.com/EichlerLab/pav).

MEIGA-PAV (https://github.com/MEIGA-tk/MEIGA-PAV)

tiSNP detection (https://github.com/Hufsah-Ashraf/ti-SNPs_detection)

Detection of altered SD organization (https://github.com/WHops/mcnv-inv)

## Data availability

Additional materials, including supplementary tables and figures, are not part of this preprint and will be released ensuing formal peer-review. All data generated has been made publicly available via the International Genome Sample Resource (IGSR); see www.internationalgenome.org/data-portal/data-collection/hgsvc2. The full inversion callset is available in **Table S2**. Raw genomic datasets are available at the International Nucleotide Sequence Database Collaboration (INSDC) under the following accessions and project IDs: Illumina WGS (PRJEB37677), RNA-seq (ERP123231), Bionano Genomics (ERP124807), PacBio (PRJEB36100), and Strand-seq (PRJEB12849, PRJEB39750). Cell lines/DNA samples were obtained from the NIGMS Human Genetic Cell Repository at the Coriell Institute for Medical Research. All phased assemblies used in this study were obtained from their original resources as described in Ebert et al. 2021 (PGAS v12 assemblies) and in Ebler et al. 2020 (PGAS v13 hifiasm assemblies). TEMPORARY SOURCE: since the PanGenie paper (Ebler et al. 2020) has not yet been formally published, we state the source for the PGAS v13 hifiasm phased assemblies here in explicit form 10.5281/zenodo.5119259. Publicly available data (PacBio and ONT) used in this study are reported in **Table S13**.

## Notes

### Competing Interest Statement

The authors have declared no competing interest.

